# Form follows function: Variable microtubule architecture in the malaria parasite

**DOI:** 10.1101/2022.04.13.488170

**Authors:** Josie L Ferreira, Vojtěch Pražák, Daven Vasishtan, Marc Siggel, Franziska Hentzschel, Emma Pietsch, Jan Kosinski, Friedrich Frischknecht, Tim W. Gilberger, Kay Grünewald

## Abstract

The malaria parasite undergoes a series of extensive morphological changes within its human host and mosquito vector. A scaffold of microtubules beneath a peripheral double membrane establishes and maintains the distinct shapes of all infectious forms, but the underlying structural basis remains unknown. Here we applied *in situ* electron cryo-tomography after focused ion beam milling to follow changes in the microtubule cytoskeleton throughout the *Plasmodium* life cycle. This revealed an unexpected level of structural and architectural diversity so far not observed in other organisms. Microtubules in migrating mosquito forms consist of 13 protofilaments reinforced by interrupted luminal helices. Conversely, gametocyte microtubules consist of 13 to 18 protofilaments with doublets, triplets and quadruplets of varying arrangements. We show the microtubule cytoskeleton within the native cellular context, highlighting structurally diverse apical rings which act as microtubule organising centres. This provides a unique view into a relevant human pathogen with an unusual microtubule cytoskeleton.

## Introduction

The eukaryotic protozoan parasites of *Plasmodium spp*., the causative agents of malaria, have a complex life cycle alternating between mosquito vector and vertebrate host. Extensive coevolution with the two hosts has resulted in the pathogen utilising a multitude of highly specialised and morphologically distinct cell types, here referred to as forms. Malaria remains one of the most dangerous and persistent infectious diseases world-wide causing 627 000 deaths in 2020 (WHO, 2021) with *P. falciparum* being the most lethal of the five human-adapted species. Occupying multiple distinct niches within the human host and mosquito vector, *P. falciparum* undergoes substantial structural and morphological changes, metamorphosing between each specialised form (Fig. 1A).

**Figure 1:**
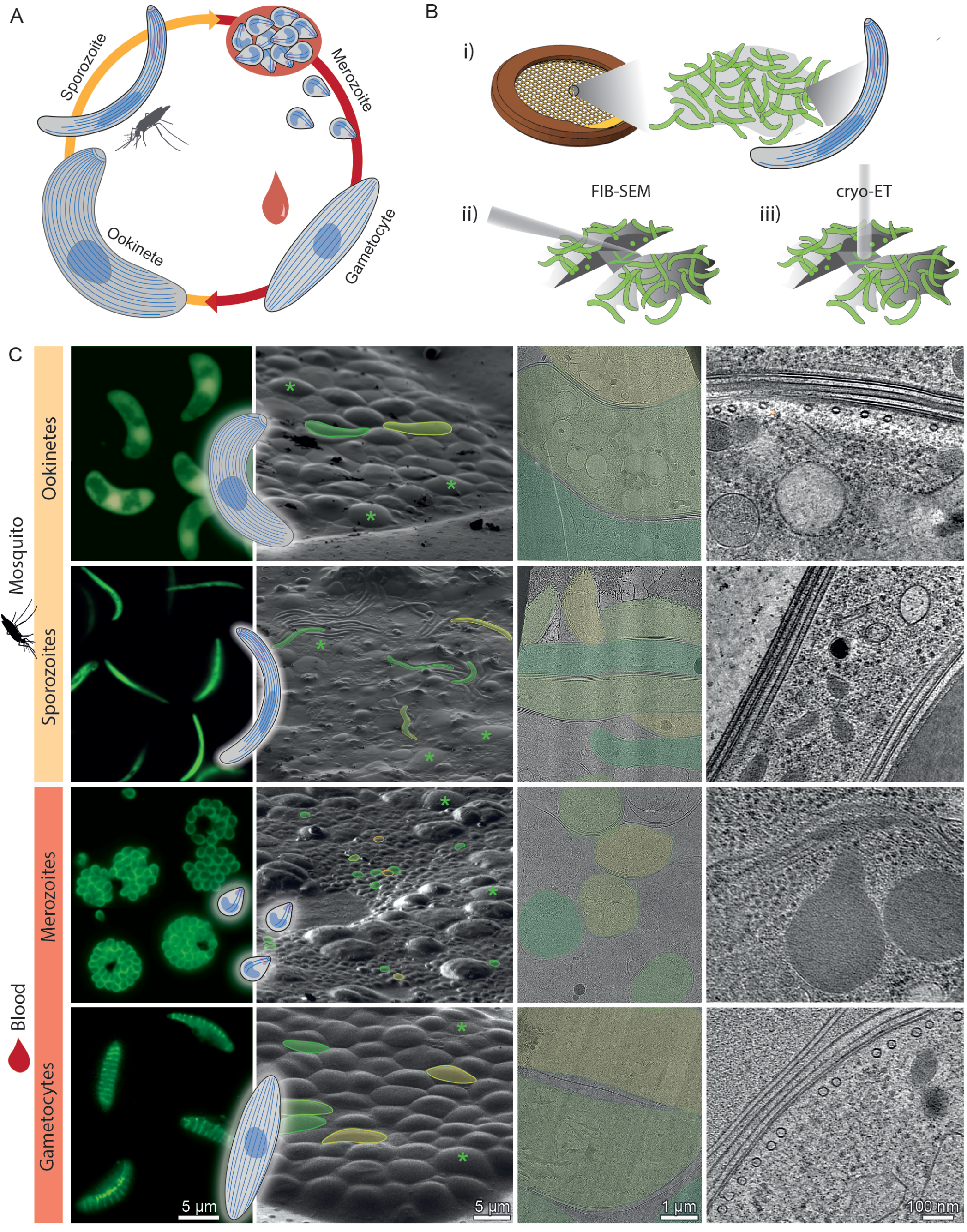
Imaging parasites across the *Plasmodium* life cycle: from live parasites to high resolution 3D volumes. **A**. Simplified *Plasmodium* life cycle of parasite forms studied here. Sporozoites are injected into the host. After differentiating in the liver, merozoites are released into the blood. The majority enter an asexual replication cycle (merozoites) and a small percentage commit to becoming “pre-sexual” gametocytes. Gametocytes are taken up by a mosquito and after fusion of male and female gametes in the mosquito gut, zygotes transform into ookinetes. Ookinetes cross the mosquito midgut, develop into oocysts and form thousands of sporozoites, which migrate to the salivary glands. **B**. Schematic representation of our workflow: i) live parasites are vitrified on EM grids. ii) cells are thinned into “lamella” and then iii) imaged by transmission electron microscopy (TEM). Tilt-series are collected and computationally reconstructed into 3D volumes. **C**. Columns 1-4: images of parasites at different workflow steps. 1: Compositions of fluorescence images of cells highlighting overall parasite shape. Inset: Cartoon representation of each stage. 2: Scanning Electron Microscopy (SEM) micrographs showing *Plasmodium* parasites (some highlighted in yellow and green) surrounded by host cells (green asterisks). 3: TEM micrographs showing overviews of lamellae. 4: Slices through example tomograms.

The pellicle is a common structural feature found underneath the plasma membrane of apicomplexan parasites, including the genera *Plasmodium* and *Toxoplasma*. Its two main constituents are a network of subpellicular microtubules (SPMTs) underneath a double membrane known as the inner membrane complex (IMC). Rather than being a static feature, the pellicle is broken down and rebuilt during the maturation of each stage and is accompanied by a different set of associated proteins (Ferreira et al., 2021; Harding and Frischknecht, 2020). This *de novo* re-organisation is a driving force behind the parasite’s substantial morphological changes and SPMTs play a key role in this process (Harding and Meissner, 2014). SPMTs and their interactions with the IMC provide the scaffold for stage-specific cellular architecture and are essential for motility and tissue penetration.

Microtubules are a common cytoskeletal component in all branches of eukaryotes. They form the tracks for intracellular transport, structural support for locomotion and spindles for chromosome segregation. The polymerisation of α-/β-tubulin heterodimers leads to the formation of protofilaments which interact laterally to form hollow cylinders of 13-protofilaments: the canonical microtubule. The resulting arrangement of tubulin heterodimers is pseudo-helical. Accordingly, the term “seam” is used to refer to the break in helical symmetry between protofilament 1 and 13. As a rule, cytoplasmic microtubules are composed of a single tube (singlet microtubule). Doublets (typically consisting of 13+10 protofilaments) and triplets are found in cilia and flagella (Fisch and Dupuis-Williams, 2011; Leung et al., 2021). The 13-protofilament microtubule is considered canonical due to its prevalence within cells of all eukaryotic subgroups (Chaaban and Brouhard, 2017). However, microtubules assembled from purified tubulin (commonly from bovine or porcine brain) *in vitro* form 9- to 16-protofilament microtubules, with 14 being the most frequent (Chaaban and Brouhard, 2017; Pierson et al., 1978). As these polymorphisms are not normally found in cells, stipulation of a uniformly 13-protofilament microtubule population therefore needs to be established during nucleation by external factors. There are four known mechanisms of controlling the number of protofilaments in a microtubule: Templating via the γ-tubulin ring complex (γTuRC) (Oakley et al., 2015; Wieczorek et al., 2020), setting a specific inter-protofilament angle through lateral binding of proteins such as doublecortin (Manka and Moores, 2020; Moores et al., 2004), expression of different tubulin isoforms (Fukushige et al., 1999) or post-translational modifications (Cueva et al., 2012).

In metazoans, nucleation of microtubules usually occurs at discrete locations, organised by a microtubule organising centre (MTOC). In the invasive forms of apicomplexan parasites, a unique MTOC at the apical end, the Apical Polar Rings (APR), coordinate SPMTs (Russell and Burns, 1984). APRs form a prominent part of the apical assembly which defines the parasite polarity. A feature unique to apicomplexan tubulin is its formation of L-shaped “semi tubules” which, with accessory proteins, form a helical structure termed the conoid (Hu et al., 2002; Sun et al., 2022), located within APRs. The conoid is a specialised structure involved in invasion. Although only characterised in *Toxoplasma*, homologues of some conoid components are expressed in *Plasmodium spp*. (Koreny et al., 2021).

Eukaryotic tubulins are highly conserved, but there are appreciable differences that make apicomplexan microtubules stand out. *Plasmodium sp*. express one β- and two α-tubulin isoforms; the two α-tubulin isoforms share ∼95% sequence identity. The parasite’s SPMTs are extremely stable, resistant to most classical depolymerisation treatments such as cold treatments (Liu et al., 2016), addition of microtubule depolymerising drugs (Morrissette and Sibley, 2002) and detergents (Cyrklaff et al., 2007; Tran et al., 2012). Due to the difficulty in obtaining purified tubulin from *Plasmodium* parasites (Hirst et al., 2022), much of our understanding of *Plasmodium* microtubules formed over the last decades has been inferred from other eukaryotic organisms. For example, reconstituted metazoan microtubules have been used to solve structures of *Plasmodium* microtubule associated proteins (MAPs) and motor proteins (Cook et al., 2021). As no plasmodium tubulin structure is available, metazoan or plant tubulin is used as a template for modelling and docking to study potential antimalarial therapeutics (Chakrabarti et al., 2013; Soleilhac et al., 2018).

Recently, expansion microscopy has provided overviews of the SPMT scaffold in some *Plasmodium* species and forms (Bertiaux et al., 2021; Rashpa and Brochet, 2022) and results from the first purification of tubulin from asexual blood forms were published (Hirst et al., 2022). Even with these advances, our understanding of *P. falciparum’s* microtubules remains incomplete, particularly in less-studied forms such as the mosquito forms and “pre-sexual” gametocytes. Our current hypotheses on microtubule control, organisation and trafficking in *Plasmodium* are based on the assumption that *Plasmodium* microtubules are canonical. There is also a need to integrate the results of *in vitro* studies in the context of the native parasite cell biology by more holistic approaches. *In situ* electron cryo-tomography (cryo-ET) is the technique of choice for this purpose. It is uniquely suited to bridge the gap between cell and structural biology, and to generate new observations and hypotheses. Cryo-ET can be used to generate 3D volumes (tomograms) of otherwise unperturbed, unstained whole cells in their native environment. Used together with sub-volume averaging (SVA), the increased local resolution for repeating structures within tomograms can be used to generate pseudo-atomic models and to analyse interrelationships between molecules. Focused ion beam (FIB) milling facilitates cryo-ET imaging of specimens that were previously unsuitable for cryo-ET due to their thickness, which includes many of the *Plasmodium* forms.

In this study, we used FIB-milling, cryo-ET and SVA (Fig. 1) to analyse the structural diversity of microtubules in *Plasmodium spp*. Focusing primarily on SPMTs, we show an unexpected level of structural variation between the individual forms. Assembled in the 3D context of the different parasite cells, our *in situ* data provides insights into the organisation and clues about the templating of microtubules throughout the *Plasmodium* parasite’s life cycle.

## Results

### Visualisation of native microtubules throughout the *Plasmodium* life cycle by *in situ* electron cryo-tomography

To determine the *in situ* structures of microtubules in different *Plasmodium* forms, we performed cryo-ET on two mosquito forms (sporozoites and ookinetes), and two human forms (merozoites and gametocytes, Fig. 1). *P. falciparum* sporozoites were obtained from mosquito salivary glands, *P. falciparum* merozoites and gametocytes from cultured blood. As *P. falciparum* ookinetes were unobtainable in sufficient quantities, ookinetes from the rodent-infecting *P. berghei* were obtained from *in vitro* differentiated gametocytes. We performed targeted FIB milling (Fig. 1B) to produce lamella with thicknesses between 50 nm and 300 nm at locations containing parasites. Initially, correlative light and electron microscopy (CLEM) was used to identify parasites vitrified on EM grids. We later found that their distinctive shapes in the SEM (Fig. 1C, column 2) allowed targeting without correlation. For each stage we acquired between 50 and 100 tomograms (roughly corresponding to the equivalent number of individual cells) from ∼20 lamellae, resulting in good coverage of subcellular regions. The distinct shape of microtubules allowed us to locate and apply SVA to all microtubules in the datasets which, in some cases, resulted in complete models of microtubule apical organisation (Fig. S1). To avoid potential bias, all ∼850 individual microtubules in the dataset were analysed independently with SVA. This allowed us to investigate structural variability, including the number of protofilaments and relative polarity, both within each cell and between the four life cycle forms.

### Sporozoite and ookinete microtubules consist of 13-protofilament microtubules with an interrupted luminal helix

Sporozoite and ookinete cells are elongated with many SPMTs lining nearly their entire length. Both move by gliding and can penetrate host tissues. Sporozoites are faster and have a smaller cell cross section (1 µm diameter, Fig. 1, compared to ∼4 µm of ookinetes). While ookinetes have around 60 microtubules, sporozoites typically have 11-16 microtubules, depending on the *Plasmodium* species.

At first glance, the lumen of SPMTs in both sporozoites and ookinetes contained a (pseudo-) helical density with ∼ 8 nm periodicity, consistent with previous predictions based on Fourier analysis of *P. berghei* sporozoite microtubules (Fig. 2A, 3) (Cyrklaff et al., 2007). Analogous structures, named Interrupted Luminal Helices (ILH) were first observed in flagellar ends of human spermatozoa (Zabeo et al., 2018), and more recently in equine and porcine spermatozoa (Leung et al., 2021), and tachyzoites of *Toxoplasma gondii* (Sun et al., 2022; Wang et al., 2021). We adopted the nomenclature of Zabeo et al. To generate an unbiased EM density map, 422 sporozoite and 177 ookinete SPMTs were analysed by SVA without applying any (pseudo-) symmetry. Both of the resulting structures showed 13-protofilament microtubules with twice-interrupted luminal helices (Fig. 2B-D, 3C). They were similar to the recent *T. gondii* SPMT structures determined in detergent solubilised cells (Sun et al., 2022; Wang et al., 2021). *T. gondii* ILH consists of thioredoxin-like proteins 1 and 2 (TrxL1, TrxL2) and subpellicular microtubule protein 1 (SPM1). *Plasmodium. spp* have homologues of TrxL1 (PfTrxL1) and SPM1 (PfSPM1) but lack TrxL2, which fills in one of the two “breaks” in the *Toxoplasma* ILH (Fig. S2). Consistently, we exclusively observed twice-interrupted luminal helices. PfTrxL1 and TgTrxL1 share 61% sequence identity (Fig. S2D), PfSPM1 and TgSPM1 are highly conserved (Fig S2E) (Tran et al., 2012), and the model of *T. gondii* SPMT assembly (pdb 7MIZ) fits well into our EM maps (Fig. 2C, D). We are therefore confident that *P. falciparum* ILH consists of 10 copies of PfTrxL1, likely with an equivalent number of PfSPM1 separated into two half-crescents (Fig. 2C).

**Figure 2:**
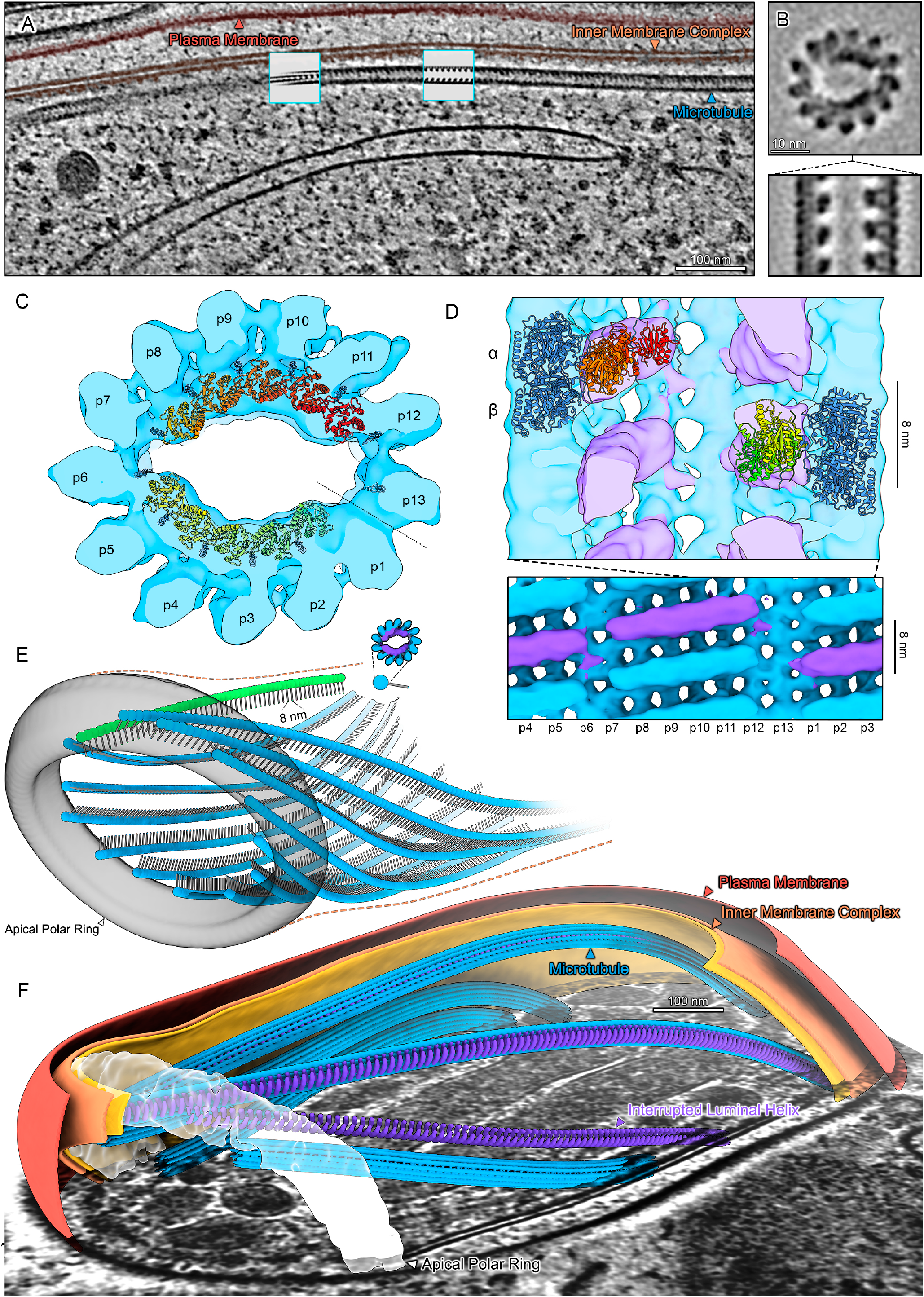
Sporozoite subpellicular microtubules (SPMTs) contain an interrupted luminal periodic density inside 13 protofilament microtubules. **A**. Slice through a tomogram illustrating the overall architecture of the pellicle within a cell. One SPMT is seen weaving in and out of the slicing plane. The luminal striations are clear in the SPMT. Insets: Slices through EM maps (shown in B, C, D) placed back into the tomogram at positions determined by SVA. **B**. Orthogonal slices through the EM map. **C**. Isosurface representation of the EM map with TrxL1 and SPM1 fitted into the EM density. p1-p13 = protofilament numbers, dotted line = seam position. **D**. Top: Section through the EM map showing the position of α/β tubulin dimer relative to TrxL1. Bottom: Radial projection with one period of the ILH highlighted in purple. **E**. The apical pole of a *P. falciparum* sporozoite with a full set of microtubules (13 + 1, coloured blue and green, respectively). The APR is represented by an isosurface, the SPMTs as “pin models” where the pinhead marks the centre and the point is oriented towards the seam. **F**. Segmented sporozoite apical pole. The tubulin density of two SPMTs was hidden to reveal the ILH.

**Figure 3:**
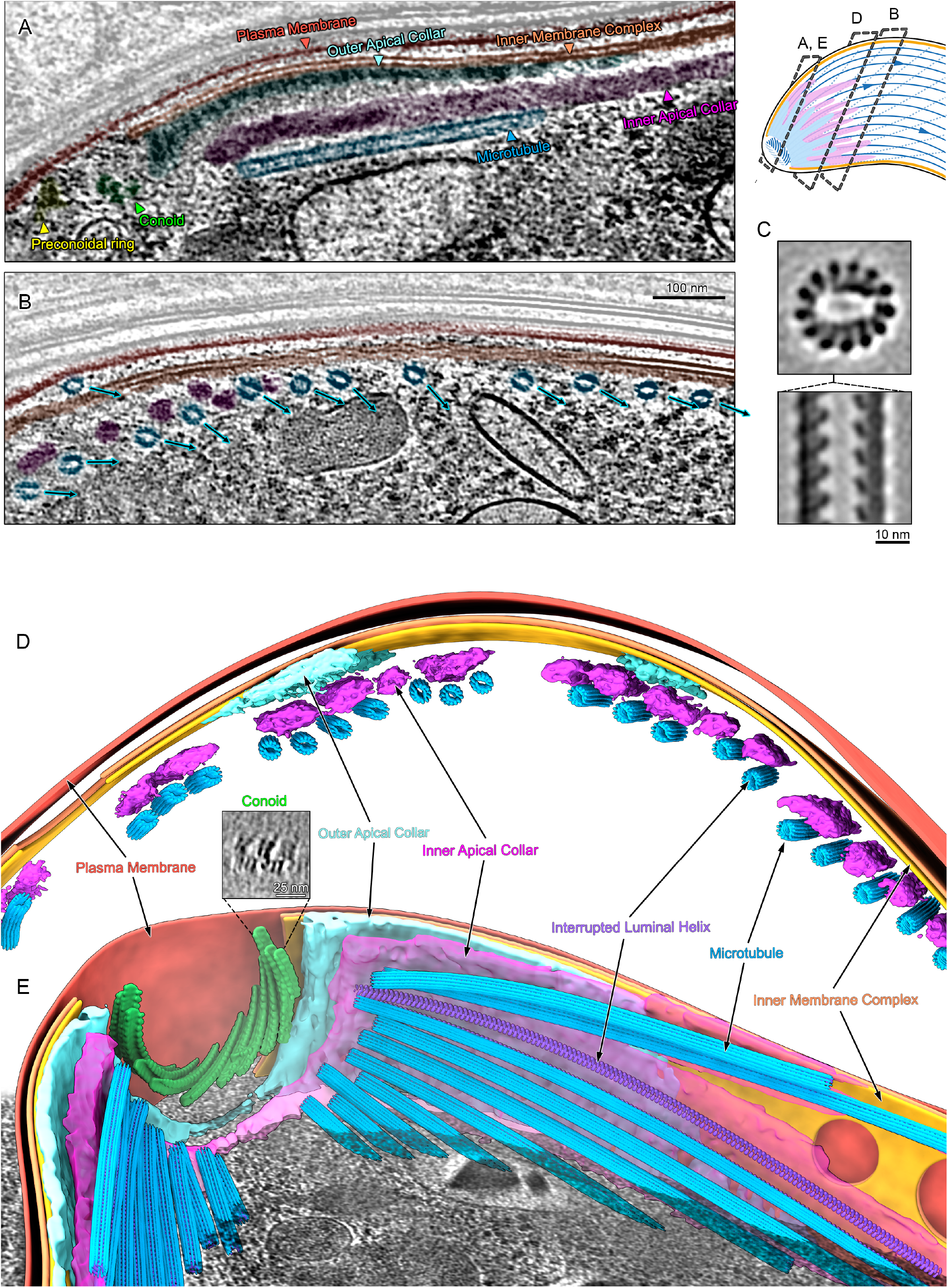
Ookinete SPMTs contain interrupted luminal helices, but are organised in a structurally different apical pole. **A**. Tomogram slice through the apical end of an ookinete with SPMTs running longitudinally. A cartoon on the right shows approximate centre positions of lamella for A, B and D. **B**. Slice through an ookinete apex with SPMTs cut transversally, the annotation colours are the same as in A. SPMTs get closer to the IMC as inner apical collar tapers off (from left to right). The rotational orientation of SPMTs (centre to seam) is indicated with arrows. **C**. Orthogonal sections through EM map determined by SVA of ookinete SPMTs. **D**. Segmentation of a tomogram near ookinete apex with transversally cut SPMTs, highlighting the two apical collar layers between the SPMTs and IMC. **E**. Segmentation of the apical end of an ookinete. The rotational axis of the apical collar is aligned with the parasite apico-basal axis. The outer apical collar is in direct contact with the IMC and has few, wide tentacles which extend ∼ 0.6 µm in the basal direction. The inner apical collar is in contact with the staggered SPMT minus ends, and has thinner tentacles which extend ∼ 1 µm in the basal direction. The tubulin density of one SPMT was hidden to reveal the ILH. Inset shows a slice through an average volume of the conoid periodic structure.

Although most microtubule structures solved to date are close to circular in cross section, microtubules with ILH are flattened along an imaginary axis roughly between protofilaments 2 and 9. This implies that the ILH stabilises the inter-protofilament angle at ∼ 26°, deviating by ∼ 2° from the theoretical 27.7° of canonical 13-protofilament microtubules (Fig. S2A, B). The deviation from circular cross-section accumulates over 5 subunits and is then compensated for by 37° bending between protofilaments 6 and 7, and 12 and 13.

### SPMTs are coordinated at structurally diverse ookinete and sporozoite apical poles

In invasive parasite forms, the APR acts as a unique MTOC, coordinating assembly of SPMTs. Consistent with this, all SPMTs in both mosquito forms had the same polarity, with the minus ends at the apical pole. The minus ends were blunt and without a γTuRC cap, although there may be additional large MAPs in the lumen of ookinete minus ends (Fig S3A). Despite both forms containing 13-protofilament SPMTs lined with ILH, there were large scale differences in architecture of the apical poles of these two forms (Figs. 2, 3).

In sporozoites, the IMC terminated with an apparently amorphous protein ring - the APR (Fig. 2E, F, Fig. S1). In two tomograms we could see the full APR with a set of regularly interspaced 13+1 SPMTs. SPMT minus ends were in an apparent direct contact (Fig. S1B) and flush with the apical edge of the APR. However, each SPMT had a different “angle of attack” relative to the ring symmetry axis which is tilted by roughly 45° from the parasite apico-basal axis (Fig. S1C, D, (Kudryashev et al., 2012). The SPMT-APR contact, therefore, needs to allow a high degree of flexibility. In contrast, the SPMT radial orientation was constrained, with protofilaments 6 to 9 preferentially making contact with the APR (Fig. S1C).

Though ookinete SPMTs had radial orientation equivalent to those in sporozoites, their higher order organisation was otherwise substantially different (Fig. 3). The most obvious difference is that ookinetes had two concentric layers of amorphous protein at the apical end of the IMC, rather than a single ring (Fig. 3A, D, E). Due to the difference in morphology, we therefore favour the term “apical collar” (AC) (Koreny et al., 2021), but APR is equally valid due to functional similarity. The apical rim of both AC layers was flat, but separated into “tentacles’’ on their basal side (Fig. 3B, D, E). The apical rim of the inner AC was in direct contact with the SPMT minus ends. These originated at staggered positions and with tighter inter-SPMT spacing than in sporozoites (Fig. 3E). This could be the consequence of the large number of ookinete SPMTs being constricted into a narrow ring.

### Ookinetes have a conoid consisting of L-shaped tubulin

At the apical end of the ookinete AC, we observed a structure consistent with a classical conoid (Fig. 3E). The average volume cross section is consistent with the L-shaped tubulin-based conoid described in *T. gondii* (Sun et al., 2022). The ookinete conoid measured 70 nm in height and 300 nm in diameter, resembling a retracted state. Until recently, *Plasmodium* (belonging to the class *Aconoidasida*, meaning “conoid-less”) was thought to not include a conoid. However, components of the conoid were shown to be present in the genome, to localise to the apical pole by fluorescence microscopy and structures that could correspond to a conoid were identified by classical EM in *Plasmodium spp*. (Koreny et al., 2021).

### The SPMTs and spindle microtubules of merozoites are canonical 13-protofilament microtubules

From our mosquito form data and the recent structures from *T. gondii*, it seemed plausible that SPMTs of all *Apicomplexa* contain ILH. To verify that this is a universal feature in all life cycle form, we analysed *P. falciparum* forms from the human host. The asexual blood stage, the merozoite, which egresses from a mature schizont, is a small cell with only two to three SPMTs (Fig. 4). Merozoites are short-lived; to avoid imaging non-viable cells, we stalled schizonts prior to egress using a well-characterised protease inhibitor (E64) (Salmon et al., 2001), resulting in merozoites in erythrocyte membrane “sacks”. Intending to interrogate potential differences between SPMT and spindle microtubules, we imaged at two timepoints of schizont development. For SPMTs, we imaged fully segmented schizonts (merozoites), while for spindle microtubules we aimed for dividing schizonts.

**Figure 4.**
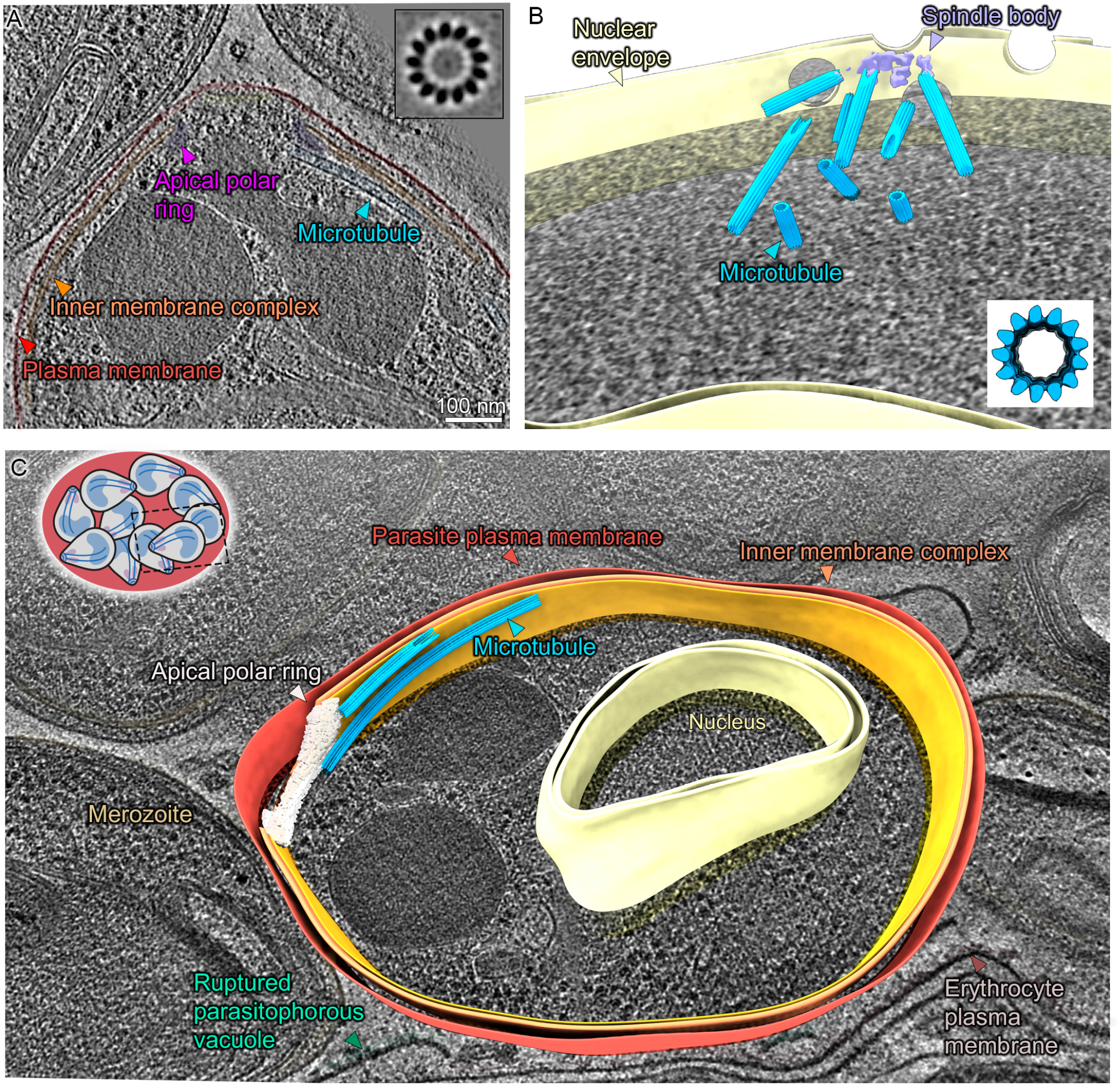
Merozoites have canonical subpellicular (SPMTs) and nuclear spindle microtubules. **A**. Slice through a tomogram of an apical pole. Inset: Slice through the SPMT EM map showing 13 protofilaments. **B**. Segmentation of a nucleus in a dividing schizont with a partial spindle body. A cluster of nuclear pore complexes are seen in proximity of the spindle. Inset: Slice through an isosurface of the SPMT EM map. **C**. Segmentation of a single merozoite within a fully segmented schizont. Note: for simplification the IMC is segmented as a continuous double membrane although multiple discontinuities were observed.

Unexpectedly, and in contrast to mosquito forms, there was no ILH in merozoite SPMTs (Fig. 4A, C). Detailed analysis by SVA showed that they were canonical, i.e. made of 13 protofilaments. Although they lack an ILH, one similarity to the mosquito forms is that merozoites had an APR to which SPMTs were bound directly. Consistent with the APR being an MTOC as in the mosquito forms, all SPMTs in merozoites had the same polarity with the plus termini facing away from the apical polar ring.

The structure of the asexual form microtubules is independent of the cellular compartment: In the nucleus, spindle microtubules radiating from a spindle pole body (Fig. 4B) were canonical and lacked an ILH. However, in contrast to the SPMT, where minus ends lacked a capping density, spindle microtubules had a clear cap consistent with γTuRC (Fig. S3C). Hence, merozoite microtubules are canonical and distinct from those in mosquito forms. This was the first indication that microtubule nucleation may be parasite-form and compartment-dependent.

### Gametocytes have non-canonical 13- to 18-protofilament singlets, doublets and triplets

The *P. falciparum* gametocyte develops from a round, then an elongated, and eventually falciform shape through five morphological stages. SPMTs start polymerising in stage II and by stage IV they completely surround the cytoplasm. We enriched gametocytes from culture at different stages in the maturation process and investigated both SPMTs and nuclear spindles to complement observations made in merozoites.

In all gametocyte stages, SPMT cross sections had noticeably large and variable diameters and consisted of a mixture of singlets, doublets, triplets (often with unusual geometries) and even a quadruplet (Fig. 5A, S4). As with the merozoite SPMTs, we found no evidence of an ILH in these “giant” microtubules. SVA was performed independently on each of the ∼170 singlet and ∼30 A-tubules of doublet SPMTs, and amazingly, showed that they consisted of 13, 14, 15, 16, 17 or 18 protofilaments (Fig. 5C, table S1). Consequently, the singlet SPMT diameters ranged from 27 to 38 nm (Fig. 5E) but were not organised with any appreciable pattern or clustering. Although some canonical microtubules were present, these only accounted for 9% of the population, while the majority of singlet microtubules had 17-protofilaments (40%) (Fig. 3C). Doublets were also “giant” and their A-tubules had a similar distribution, with 17 protofilaments being the most common (Fig. S4B). Surprisingly, considering the ubiquity of a 13-protofilament A-tubule in other organisms, there were no A tubules with 13 protofilaments.

**Figure 5.**
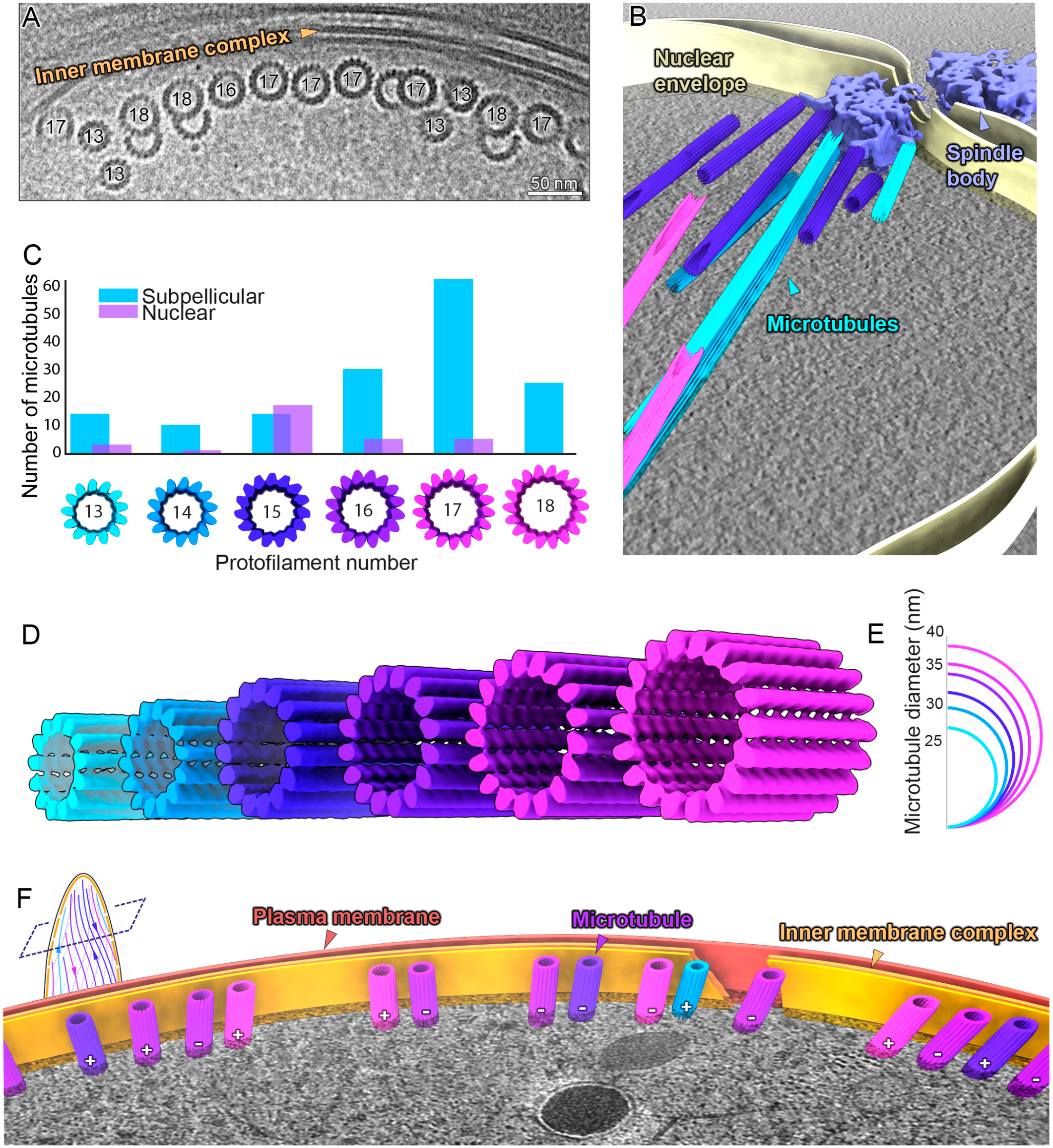
Gametocyte subpellicular microtubules (SPMTs) have a wide range of protofilament numbers with random polarity. **A**. TEM micrograph showing a row of singlet and doublet SPMTs highlighting the range of sizes. Numbers correspond to protofilament numbers (of the A tubule in doublets). Micrographs at two different tilt angles were stitched to show SPMT transversal views. **B**. Segmentation of a stage III gametocyte nucleus with a spindle body at a nuclear pore complex. Microtubule colours correspond to protofilament numbers as shown in C,D. **C**. Histogram of the distribution of different protofilament numbers in SPMTs (blue) N=155, and nuclear spindles (lilac) N=31. Distributions are significantly different (p = 5 × 10^−8^). **D**. Isosurfaces of microtubules from SVA with protofilament numbers from 13 to 18. **E**. Schematic representation of the differences in microtubule diameter. **F**. Segmentation of a stage III gametocyte with transversely sectioned SPMTs. Microtubule colours correspond to protofilament numbers as shown in C,D. ‘+/-’ annotates the polarity of each microtubule.

Gametocyte cells do not have clear cellular polarity, unlike the distinctly polar cells of the three invasive forms described above (Figs. 2-4). Although recent expansion microscopy data hints at a presence of ring-like structures at gametocyte extremities (Rashpa and Brochet, 2022), there was no evidence of an APR-like structure that could act as a MTOC at gametocyte poles. SPMTs originated at apparently uncoordinated positions and there was no discernible pattern in the distribution of SPMTs along the IMC with respect to the number of protofilaments or secondary tubules. (Fig. 5F, table S1). Considering that SPMTs in the three earlier forms lacked a γTuRC cap, we had initially hypothesised that SPMT nucleation factors may be physically associated with the APR or AC. However, focused analysis of the gametocyte minus ends showed that they were also uncapped (Fig. S3). Clearly, the diversity in protofilament numbers sets the gametocyte SPMT population apart from other forms. Whether this is due to a different nucleation mechanism or a different tubulin composition could be gathered by investigating microtubules outside of the pellicle.

### Gametocyte non-canonical microtubules are independent of cellular compartment

Other than SPMTs, there are two more microtubule populations in gametocytes: nuclear spindle (or hemi-spindle (Sinden et al., 1978)) and cytoplasmic. Cytoplasmic microtubules are short-lived and only appear in early stage gametocytes on the opposite side of the cell from the developing pellicle (Schneider et al., 2017). Nuclear microtubules were seen frequently in stage III/IV parasites, and a full spindle pole body was present in one tomogram (Fig. 5C).

The large range in protofilament numbers in gametocytes is independent of the cellular compartment. Although cytoplasmic microtubules were rare, we observed a mixture of protofilament numbers (13 to 16, Fig. S4E) and while there was an indication of clustering by protofilament size, this could not be confirmed due to low numbers. In the nucleus, microtubules ranged from 13 to 17 protofilaments, with 15 being most common (Fig. 5B, C). This was unexpected as their negative ends were clearly capped with a structure consistent with γTuRC irrespective of the microtubule diameter and protofilament number (Fig. S3D). The distribution of protofilament numbers in nuclear microtubules was significantly different from SPMTs, but the wide range was puzzling considering that they were γTuRC nucleated. Together, these data support the hypothesis that microtubule nucleation is parasite form and cellular compartment dependent.

### SPMT to IMC distance is conserved in all parasite forms pointing to a universal linker protein

Due to the nature of the data being acquired *in situ*, we were able to quantify the SPMT architecture with respect to the IMC. To elucidate whether the SPMT-IMC interaction is mediated in the same manner in all parasite forms, we measured the shortest distance (dIMC) from the surface of each SPMT to the inner IMC membrane. A similar dIMC in all parasite forms would imply that the same proteins are involved in tethering SPMTs to the IMC. IMC surface coordinates and SPMT positions were determined by segmentation and SVA. Excluding the apical assemblies, there was no significant difference in the dIMC between the four parasite forms (with a common median of 18 nm and standard deviation of 10 nm, Fig. 6D). This distance increased at the ookinete apex to ∼ 50 and ∼100 nm in order to accommodate the AC, but the distance from the SPMT surface to the closest surface, the inner AC, was comparable to dIMC (∼13 nm, Fig. 6A). Thus, it is likely that there is a conserved protein, linking the SPMTs to the IMC in all parasite forms analysed.

**Figure 6:**
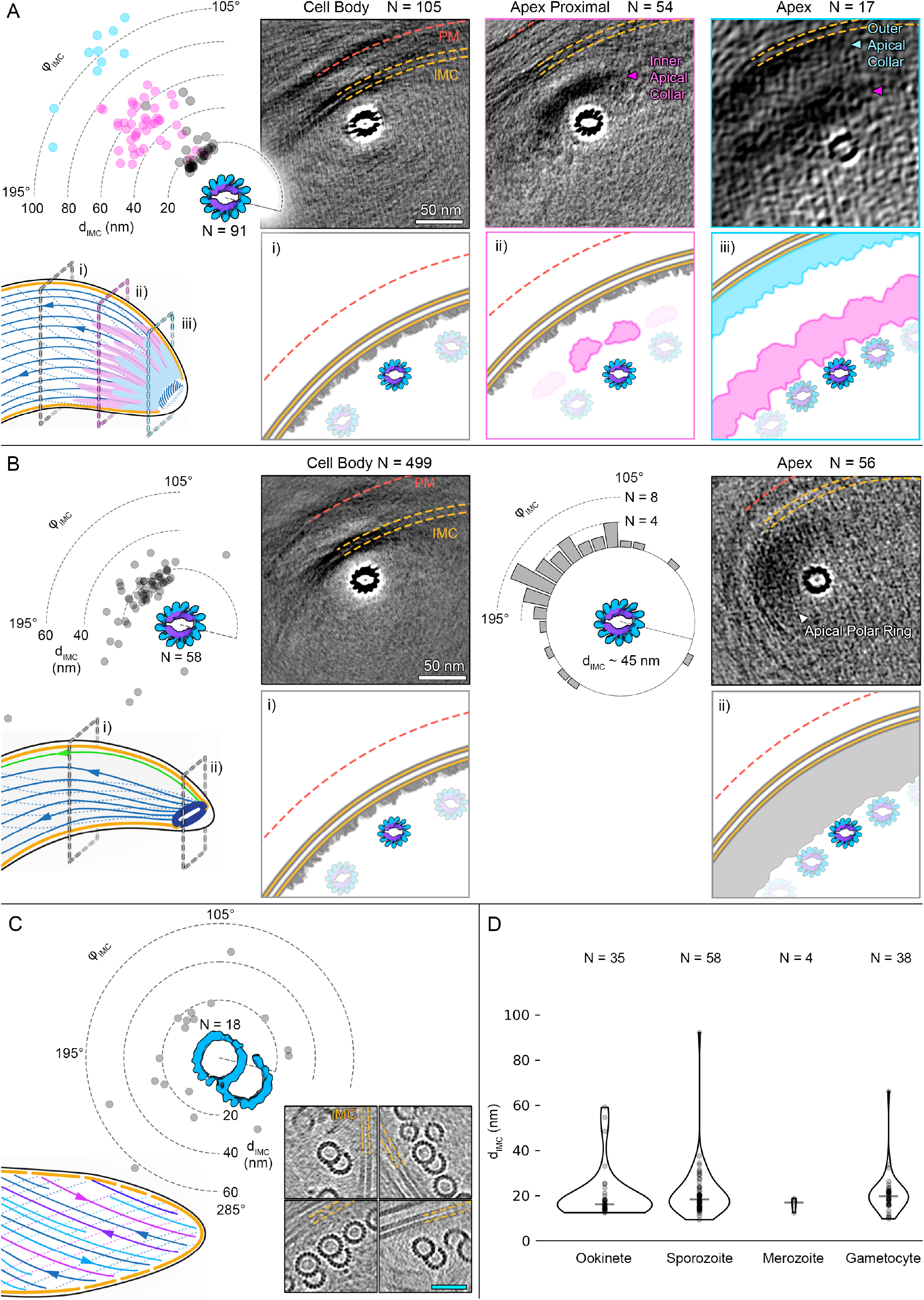
Microtubule distance from the IMC is consistent in all forms, but the interrupted luminal helix is required for setting their radial polarity. This figure shows radial scatter plots of IMC distance and orientation with respect to individual SPMTs (d_IMC_ and Φ_IMC_). Each point is the median of a single SPMT. Only SPMTs close to clearly discernible IMC could be analysed. Grayscale images in A and B are 40 nm thick slices (average projections) through EM maps of SPMTs determined at different parts of the cell (indicated by cartoons on the left-hand side of each panel). As a result of the consistent distance and orientation, IMC and APR components can be seen in the SPMT EM maps after extracting large subvolumes (200 nm edges). Number of microtubules in each average volume is indicated. Schematic representations are below each section. **A**. Ookinete (4 data points were excluded for visualisation purposes, full set in Fig. S5A). **B**. Sporozoite. The IMC wraps around the torus-shaped APR, which means that d_IMC_ and Φ_IMC_ could not be measured in the same manner. Instead, Φ_IMC_ was determined to the closest point on the APR and represented as a histogram. **C**. Gametocyte. Grayscale images are slices through individual tomograms showing different orientations of doublets. **D**. Violin plot comparing d_IMC_ between forms, widths are scaled according to the amount of data.

**Fig. 7:**
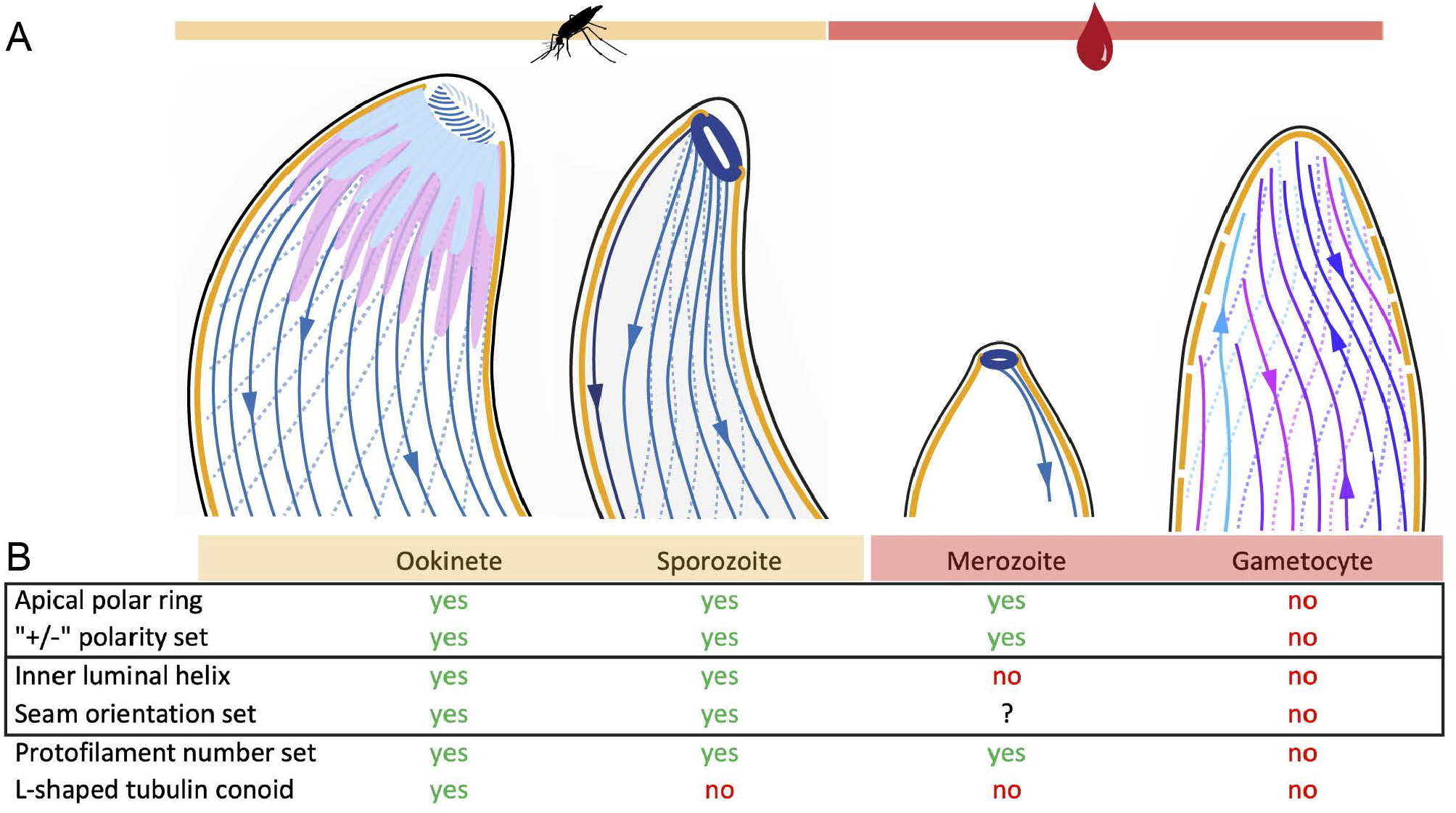
Structural diversity of *Plasmodium* microtubules across the life cycle. **A**. Cartoon representations of the four *Plasmodium* forms analysed. **B**. Table summarising the main architectural differences in the four forms. Solid outlines indicate properties that are likely correlated: the presence of an apical polar ring sets the SPMT polarity and ILH is required to set the seam orientation with respect to the IMC.

### The interrupted luminal helix controls the rotational orientation of SPMTs with respect to the IMC

With the hypothesis that a common linker is tethering the SPMTs to the IMC in all forms and with the knowledge that mosquito form SPMTs have a preferred radial orientation (Figs. 2E, 3B, S1B,C), we set out to look for the linker binding site. A clear candidate, due to it being a distinct asymmetric structural feature, was the seam. We measured the angle ΦIMC between the seam, the microtubule centre and the closest point on the IMC surface.

The first prerequisite for this relative angle measurement is the knowledge of the absolute position of the seam. This could be determined by SVA in ookinetes and sporozoites due to the asymmetric nature of the ILH. In gametocytes we made use of B tubules of doublets as a proxy for locating the seam, but this relied on one of two assumptions to be valid. The first is that the branch point is equivalent to that of canonical 13-protofilament doublets (i.e. between protofilaments n - 3 and n - 2, where n is the number of protofilaments in the A tubule (Li et al., 2021; Ma et al., 2019). Alternatively, the branch point could be in the same, but unknown, position relative to the seam regardless of the number of protofilaments in the A tubule. The latter assumption would invalidate the absolute values of measured angles but the relative distribution would still hold.

Analysing the mean ΦIMC for all possible SPMTs revealed that only mosquito stage SPMTs have a set radial polarity (Fig. 6A, B, S5). Surprisingly, the seam points away from the IMC with protofilament 8 oriented towards it instead. When we classified the SPMTs into specific locations on the cell body (specifically at the APR in sporozoites and the ACs in ookinetes) we saw no substantial differences in ΦIMC (Fig. 6A,B, S5). In contrast to the mosquito forms, no clustering of ΦIMC was observed in gametocytes (Fig. 6C).

These results were inconclusive with regards to a universal binding site on all SPMTs, but it suggested that the ILH was involved in setting the radial orientation. We inspected the mosquito stage SPMT EM density map for possible clues; even at low contour levels there was no linking density between the SPMT and IMC, indicating that any linker protein is flexible or present at a low occupancy. Nevertheless, there was some evidence of radially-asymmetric decoration with an unknown protein density between protofilaments 10, 11 and 12, with additional density emanating radially outwards from protofilaments 11 and 12 (Fig. S6). If these densities correspond to a part of the SPMT-IMC linker, there would likely need to be additional connections to established the observed radial orientation. Thus, although the precise details of a linker are yet to be elucidated, we present evidence for asymmetric, ILH-dependent MAP decoration.

## Discussion

### Large morphological changes between parasite forms are accompanied by extensive alterations in microtubule structure and organisation

The extensive morphological changes undergone by the *Plasmodium* parasite have been documented by light microscopy and classical EM (Cox, 2010). The pellicle was assumed to be a relatively constant motif across the different forms; we show that this similarity is limited only to low resolution features and the individual components are highly varied. In the two mosquito forms, sporozoites and ookinetes, SPMTs consist of 13 protofilaments reinforced by an ILH. The ILH is not present however, in the 13-protofilament SPMTs of blood stage merozoites, nor in gametocytes. Gametocyte SPMTs had a seemingly random number of protofilaments between 13 and 18, but most commonly 17. A subset of these were doublets, but triplets and quadruplets were observed, some with unusual architectures so far only observed *in vitro*. The apical poles across the four forms had large organisational differences, with the ookinete being the most elaborate and containing a conoid. There was one feature unifying the disparate cytoskeletal structures: the spacing between the *Plasmodium* SPMTs and the IMC is constant, hinting at the same unknown protein mediating this link.

### Interrupted Luminal Helices: More than just structural stabilisation?

Our observations contribute to a growing list of organisms that make use of an ILH in their microtubule cytoskeleton, which now includes the SPMTs of apicomplexans (Sun et al., 2022; Wang et al., 2021), and the cilia and flagella of mammals (Leung et al., 2021; Zabeo et al., 2018). With our observations it seems likely that ILH consisting of TrxL1 and SPM1 are conserved across the *Apicomplexa*. Intriguingly, in mammals, there is a homologue of TrxL1 that localises to lung cilia and sperm flagella (thioredoxin-like Txl-2 (Sadek et al., 2003)) and a homologue of SPM1 in axonemes (stabilizer of axonemal microtubules SAXO2 (Miranda-Vizuete et al., 2004). This suggests that the ILH may be a common eukaryotic trait.

The ILH has been hypothesised to limit turnover at the microtubule plus end and strengthen SPMTs in both the longitudinal (via SPM1) and transversal direction (TrxL1) (Leung et al., 2021; Zabeo et al., 2018). An SPM1-null mutant in *T. gondii* had reduced fitness (Tran et al., 2012). This was not seen in a second study although microtubules were more susceptible to chemical treatments when missing SPM1 or TrxL1(Wang et al., 2021). It is conspicuous that ILH occurs in locomotive structures, such as cilia, flagellar ends and apicomplexan gliding forms. Thus, we postulate that the ILH provides stability without sacrificing flexibility, and if breakages occur, SPM1 keeps the ends together, allowing SPMT repair or self-healing. Where solely rigidity is required, *Plasmodium* gametocytes have evolved a dense sheet of SPMTs with a large number of protofilaments, doublets and triplets.

The function of SPMTs as cytoskeletal elements relies on a physical connection to the IMC. Studying the linkage between the SPMTs and the IMC, we found that the rotational orientation of ILH-containing SPMTs with respect to the IMC was set. Surprisingly, the seam, where microtubules depart from helical symmetry, was not oriented towards the IMC. This was unanticipated as the seam is the most asymmetric feature that can be accessed by externally binding MAPs. Instead, it is the ILH that sets the rotational orientations. Gametocytes, which lack an ILH, have an apparently random radial polarity. Our evidence for this was the large variance of doublet ΦIMC, but beyond this it is also difficult to reconcile the supertwist of non-canonical microtubules with a linker having a preference for a specific protofilament. We propose that the radial polarity of 13-protofilament SPMTs in merozoites is also random, due to the supposed lack of an asymmetric feature like the ILH.

There are two conceivable mechanisms for how the ILH could be orienting SPMTs. Either a direct contact between an ILH component and an external MAP, or by forcing SPMTs into elliptical cross sections (Fig S2), where the inconsistent inter-protofilament angles could be recognised as binding sites. We observed weak densities emanating from protofilaments 11 and 12 and in the ridges between protofilaments 9 to 12 (Fig. S6). These densities are not consistent with known MAPs such as kinesin or doublecortin, but the resolution of our C1 EM density maps is not high enough to be conclusive. It is not clear whether they correspond to a putative SPMT-IMC linker, considering that they are not oriented towards the closest point on the IMC. A framework for understanding the evolution of asymmetric rotational orientation of SPMTs may be a recently proposed hypothesis that SPMTs originate from an ancient flagellum (de Leon et al., 2013; Wall et al., 2016). The orientation of ILH containing SPMTs is roughly consistent with the orientation of the A-tubule in axonemes with respect to the membrane.

### Subpellicular microtubule nucleation may be distinct from canonical γ-tubulin ring complex

The variations in microtubule structures among *Plasmodium* forms pose the obvious question; how are these different microtubules being nucleated? It has been suggested that SPMTs with ILH could be nucleated by SPM1 (Wang et al., 2021). TrxL1 could then set the protofilament number to 13 through the curvature of its oligomeric state. This seems plausible in isolation, but not in light of our new data. It does not explain how the giant microtubules of gametocytes or merozoite 13-protofilament SPMTs, which lack the ILH, are nucleated. Further, the curvature of TrxL1-bound protofilaments departs from that of a canonical 13-protofilament microtubule (Fig. S2) and there is a gap in the ILH between protofilaments 13 and 1 where more protofilaments could conceivably be inserted.

All forms with APRs (ookinetes, sporozoites and merozoites) have 13-protofilament SPMTs. We initially hypothesised that the nucleation mechanism could be linked to the APR, possibly through γTuRC or a doublecortin-like MAP setting the inter-protofilament angle. By extension, gametocytes, which lack an APR and have non-canonical microtubules (i.e. the inter-protofilament angles are variable), would utilise a different mechanism. However, the minus ends of all SPMTs, in all four parasite forms, were uncapped (Fig. S3), which due to the stability of SPMTs, suggests that nucleation is independent of γTuRC. This is in contrast to the spindle microtubules observed in gametocytes and dividing schizonts. The minus ends of nuclear microtubules were capped with a density consistent with γTuRC (Fig. S3). This is unexpected considering that spindle microtubules consisted of 13 protofilaments in early schizonts but anywhere from 13 to 17 protofilaments in gametocytes. Intriguingly, nuclear microtubules in the ciliates *Paramecium* and *Nyctotherus* were previously shown to have a range of protofilament numbers (13 to 16), although their cytosolic microtubules consisted exclusively of 13 protofilaments (Eichenlaub-Ritter and Tucker, 1984). Based on these observations, we hypothesise that microtubule nucleation is compartment and form specific, with SPMTs nucleation being γTuRC-independent.

### Have gametocyte microtubule giants lost control?

In the large range of model organisms studied to date, protofilament number is tightly controlled, but this doesn’t seem to be the case in gametocytes. Microtubules have the propensity to self-assemble *in vitro*, albeit at concentrations considerably higher than physiological. This spontaneous self-assembly results in microtubule populations with a wide distribution of protofilament numbers (9-16 for bovine brain tubulin) (Chaaban and Brouhard, 2017; Pierson et al., 1978) including doublets and triplets. However, a nucleation-free self-assembly is unlikely to be the reason for the observed giant non-canonical SPMTs in gametocytes; they polymerise on the surface of growing IMC plates, rather than spontaneously in the cytoplasm. Furthermore, the clustering of the distinct population of cytoplasmic microtubules suggests that a level of control is available.

Instead, the increased number of protofilaments could be due to altered post-translational modification or expression of the α2-tubulin isoform. α1- and α2-tubulin have ∼ 95% sequence identity but have several amino acid substitutions at key regions. For example, α2-tubulin is missing 3 amino acids at the c-terminus, often a target for posttranslational modifications and the most variable region in the sequence corresponds to a loop which may mediate intra-protofilament interactions. These small changes could ultimately result in alterations in the microtubule lattice. Although α-2 tubulin is likely expressed in all life cycle forms (López-Barragán et al., 2011; Zanghì et al., 2018), this isoform is not essential in asexual forms (Zhang et al., 2018) and can only partially replace α1-tubulin in *P. berghei* sporozoites (Spreng et al., 2019).

Speculating about the potential function of the giant SPMTs is challenging in light of *P. falciparum* being the only *Plasmodium* species that produces elongated gametocytes (Dixon and Tilley, 2021). The maturing gametocyte has decreased deformability (Tibúrcio et al., 2012) inhibiting premature release of gametocytes from sequestration sites in the bone marrow, therefore avoiding splenic clearance (Duez et al., 2015). This is potentially a selectable trait. There is only a limited number of microtubules that can fit onto the inner surface of the IMC, and 13-protofilament SPMTs may just be too flexible. The transition from 13 protofilaments to 17 or 18 protofilaments is expected to greatly increase the stiffness of the microtubule, as a change from only 13 to 15 protofilaments was shown to result in a 35% increase in stiffness (Gittes et al., 1993). Therefore, by maximising the number of microtubules as well as assembling larger, more rigid microtubules, this may allow *P. falciparum* gametocytes to mature undisturbed in their human hosts.

## Conclusions and outlook

By utilising FIB-milling and *in situ* cryo-ET we were able to visualise the native microtubule cytoskeletal architecture in its cellular context throughout the *Plasmodium* life cycle. We show unexpectedly large architectural differences between individual forms, particularly considering the large degree of microtubule structural homogeneity among eukaryotic clades. This highlights the extreme adaptations that specialised intracellular pathogens undergo as they co-evolve with their hosts. The data presented here have clear implications for the research of new microtubule-targeting therapeutics, which needs to take into account the complexity of what are currently unknown microtubule regulatory systems as well as the structural differences relative to other model systems. Finally, we provide a framework for new research avenues, from basic *Plasmodium* biology to the fundamental biology of microtubules.

## Acknowledgements

We would like to thank the CSSB Cryo-EM and ALFM facilities for their support. We thank Till Voss for the 3D7-iGP parasites, TropIQ for providing the sporozoites and Andrew Waters and Katie Hughes for sharing the PbRFP line. Thank you to Lindsay Baker for critical feedback on the results and John Heumann and Carolyn Moores for helpful discussions and advice. Thank you to the Stackoverflow community for being a seemingly bottomless repository of knowledge.

JLF: HFSP long-term postdoctoral fellowship (LT000024/2020-L) and support for materials from Research Infrastructures for the control of vector-borne diseases (Infravec2) funded by the EU’s Horizon 2020 programme (grant agreement No 731060).

MS: EMBL Interdisciplinary Postdoc Programme under Marie Curie COFUND actions. FH: German Center for Infection Research, DZIF.

FF: DFG research networks SFB 1129, SPP 2332 and grant FR 2140/10-1. TWG: KIF-002, DFG research networks SPP 2225.

KG: KIF-002, Wellcome Trust grants 107806/Z/15/Z and 209250/Z/17/ Z, BMBF grant 05K18BHA and DFG INST 152/ 772-1, 774-1, 775-1, 777-1 FUGG to the CSSB cryoEM facility.

## Author contributions

Sample preparation: JLF, FH and EP. Data collection: JLF. Tomogram reconstruction and subvolume averaging: JLF and VP. Data analysis: JLF, VP, DV and MS. Supervision: KG, TWG, FF, JK. Writing original draft: JLF, VP. Figure preparation: JLF and VP. Writing and editing: JLF, VP, DV, MS, FH, EP, KG, TWG, FF, JK.

## Materials availability

This study did not generate new unique reagents. EM Maps will be deposited on the EMDB.

## Ethics statement

Animal experiments were performed according to FELASA and GV-SOLAS guidelines and approved by the responsible German authorities (Regierungspräsidium Karlsruhe). *Plasmodium* parasites were maintained in five to eight-week-old female Swiss mice obtained from JANVIER.

## Materials and Methods

### Cell culture

Blood forms of *P. falciparum* parasites were cultured in human red blood cells (O+ or B+, Universitätsklinikum Eppendorf, Hamburg, Germany) at 5 % haematocrit in an atmosphere of 1% O2, 5% CO2, and 94% N2 at 37°C. RPMI complete medium was used which contained 0.5% Albumax II according to standard protocols (Trager and Jensen, 1997).

#### Parasite synchronisation

To obtain highly synchronous cultures, schizont forms were harvested using 60% prewarmed percoll as previously described (Rivadeneira et al., 1983). Isolated schizonts were washed once with prewarmed RPMI, resuspended in 5% uninfected red blood cells in prewarmed RPMI and cultured for a further 3 hours under standard conditions to allow parasites to egress and reinvade. Subsequently, unruptured schizonts were lysed with 5% sorbitol. After sorbitol treatment, the remaining culture contained young ring forms with a 3-hour synchronicity window.

#### *P. falciparum* schizonts

Schizont forms of tightly synchronised 3D7 parasites expressing endogenously GFP-tagged GAPM2 (glideosome-associated protein with multiple membrane spans 2 (Bullen et al., 2009; Kono et al., 2012)) were isolated from a 10-20 mL culture (5-10% parasitaemia) using 60% percoll as previously described (Rivadeneira et al., 1983). Purified schizonts were washed twice in pre-warmed RPMI, placed into 2 ml dishes and treated with 1 µM of the PKG-inhibitor compound 2 (provided by Dr. Mike Blackman, The Francis Crick Institute, UK) to prevent rupture of mature schizonts (Baker et al., 2017; Donald et al., 2006). Compound 2-treated schizonts were subsequently cultured under standard conditions and harvested after a further 6 hours using 60% percoll. Early and segmented schizonts were washed once and then resuspended in pre-warmed RPMI without Albumax or phenol red but with the addition of 1 µM E64 (Sigma) to allow the parasitophorous vacuole to rupture but prevent rupture of the red blood cell membrane. This helped us identify very late stage schizonts in the SEM. Cells were vitrified within an hour.

#### *P. falciparum* gametocytes

Gametocyte stages were generated by targeted overexpression of the sexual commitment factor GDV1 (Gametocyte development 1) using a 3D7 inducible gametocyte producer line (iGP) as previously described (Boltryk et al., 2021) overexpressing a GFP-tagged version of the suture inner membrane complex protein PF3D7_1345600 (plasmid from (Kono et al., 2012)). Expression of endogenously DD-tagged GDV1 was induced by the addition of Shield-1 to a parasite culture containing 2-3% rings. Parasites were cultured in the presence of Shield-1 for a further 48 hours to allow reinvasion. After reinvasion, sexual committed rings were adjusted to 10% parasitaemia and cultured in RPMI medium supplemented with 10% human serum (blood group AB+) for 10 days to allow gametocyte maturation. Culture medium was changed at least once per day on a 37°C heating plate. To deplete non-committed asexual forms, gametocytes were treated with 50 mM *N-*acetyl-d-glucosamine (GlcNac) for at least 4 days. Gametocyte stages II, III, IV and V were isolated at different time points during gametocyte maturation from a 20 mL culture using 60% pre-warmed percoll. Isolated gametocytes were washed twice and then resuspended in pre-warmed RPMI without albumax, serum or phenol red and kept warm until immediately prior to freezing.

#### *P. falciparum* sporozoites

*P. falciparum* sporozoites (strain: NF54-ΔPf47-5’*csp*-GFP-Luc: expressing a GFP-Luciferase fusion protein under the control of the csp promoter, genomic integration, no selection marker (Winkel et al., 2019) were prepared at TropIQ (Nijmegen, Netherlands).Gametocytes were fed to 2 days old female *Anopheles stephensi* mosquitoes. Mosquito infection was confirmed 7 days post infection by midgut dissection. At 7 days post infection, the infected mosquitoes received an extra non-infectious blood meal to boost sporozoite production. Two weeks post infection, sporozoites were isolated using salivary gland dissection and shipped to Hamburg at room temperature.

#### *P. berghei* ookinetes

Ookinetes were produced using the *P. berghei* line *Pb*RFP (REF), a *P. berghei* ANKA line that constitutively expresses RFP. Two female Swiss mice were injected intraperitoneally with 200 µl phenylhydrazine (6 mg/ml in PBS) to stimulate reticulocytosis. Two days later, the mice were infected intraperitoneally with 20*10^6^ iRBC *Pb*RFP. Mice were bled three days post infection and 500 µl blood was transferred to 10 ml ookinete medium (RPMI supplemented with 20% (v/v) FCS, 50 µg/ml hypoxanthine, and 100 µM xanthurenic acid, adjusted to pH 7.8 – 8.0) at 19 °C. After 22 h of culture, ookinete cultures were underlaid with 5 ml 55% Nycodenz/PBS and centrifuged for 25 min at 1000 rpm without brake. The interphase containing purified ookinetes was collected, washed in PBS and immediately plunge frozen for EM.

### Fluorescence microscopy

All fluorescence images were taken on a Leica D6B fluorescence microscope equipped with a Leica DFC9000 GT camera and a Leica Plan Apochromat 60x or 100×/1.4 oil objective. Microscopy of live parasites was performed by placing 5 μl of parasites onto a glass slide and covering with a cover slip. Contrast and intensities were linear adjusted to present clear parasite shapes using Fiji and cropped images were assembled into a composition of cells for Figure 1 using Adobe Illustrator CC 2021.

### Plunge freezing

3µl of enriched parasites were applied onto a freshly plasma-cleaned UltrAufoil R1.2/1.3 300 mesh EM grid (Quantifoil) in a humidity controlled facility. Excess liquid was manually back-blotted and grids were plunged into a reservoir of ethane/propane using a manual plunger. Grids were stored under liquid nitrogen until imaging.

### Cryo FIB milling

Grids were clipped into autogrids modified for FIB preparation (Schaffer et al., 2015)and loaded into either an Aquilos or an upgraded Aquilos2 cryo-FIB/SEM dual-beam microscope (Thermofisher Scientific). Overview tile sets were recorded using MAPS software (Thermofisher Scientific) before being sputter coated with a thin layer of platinum. Good sites with parasites were identified for lamella preparation before the coincident point between the electron beam and the ion beam was determined for each point by stage tilt. Prior to milling, an organometallic platinum layer was deposited onto the grids using a GIS (gas-injection-system). Lamellae were milled manually until under 300nm in a stepwise series of decreasing currents. Milling was performed at the lowest possible angles to increase lamella length in thin cells. Finally, polishing of all lamella was done at the end of the session as quickly as possible but always within 1.5 hours to limit ice contamination from water deposition on the surface of the lamellae. Before removing the samples, the grids were sputter coated with a final thin layer of platinum. Grids were stored in liquid nitrogen for a maximum of 2 weeks before imaging in the TEM.

### Cryo-EM

FIB-milled grids were rotated by 90 degrees and loaded into a Titan Krios microscope (Thermofisher) equipped with a K2 or K3 direct electron detector and (Bio-) Quantum energy filter (Gatan). Tomographic data was collected with SerialEM (Mastronarde, 1997) with the energy-selecting slit set to 20 eV. Datasets were collected using the dose-symmetric acquisition scheme at a ±65° tilt range with 3° tilt increments. For all datasets, 5-10 frames were collected and aligned on the fly using SerialEM and the total fluence was kept to under 120e^−^/Å^2^. Defoci between 3 and 8 μm underfocus were used to record the tilt series’.

### Tomogram reconstruction

Frames were aligned on the fly in SerialEM; CTF estimation, phase flipping and dose-weighting was performed in IMOD (Kremer et al., 1996). Tilt-series’ were aligned in IMOD either using patch-tracking or by using nanoparticles (likely gold or platinum) on lamella surfaces as fiducial markers. Tomograms were binned 4x and filtered in IMOD or by using Bsoft (Heymann, 2001).

### Sub-volume averaging

#### For all averages

SVA was performed in PEET (Heumann et al., 2011). Initial 3D coordinates (model points or particle coordinates) for SVA were generated by interpolating between manually traced microtubule centres in IMOD, using scripts based on TEMPy (Cragnolini et al., 2021) and using Python libraries Numpy, Scipy and Matplotlib (Harris et al., 2020; Hunter, 2007; Virtanen et al., 2020). The vectors between pairs of points were used to set initial particle Y axis orientation (microtubule pseudosymmetry axis). Each microtubule was processed separately, aligned to a unique reference (raw average) generated by averaging the respective particles with initial orientations (Fig. S7). In order to determine its polarity and the number of protofilaments, the initial rotations around the Y axis (here referred to as Φ rotation) were randomised (Fig S7, step 2). Particles from the same forms with the same number of protofilaments were then combined for further processing. The progress of SVA was monitored by inspecting “pin models” in UCSF Chimera (Fig3. E REF) where each particle’s position and orientation were represented by a pair of markers. This allowed us to e.g.: identify and remove outliers from the linear geometry. Particles were pruned if they overlapped with others, had low cross-correlation coefficient or drifted too far from their initial positions. SPMT maps were sharpened using arbitrary B-factors using Bsoft (Heymann, 2001).

#### Sporozoite and ookinete SPMTs

Having determined the relative polarity, the necessary step for combining is finding the relative Φ rotation (rotation around the symmetry axis) between individual microtubules. This could be done by aligning each individual particle to a common reference, but would result in a large number of errors due to the low signal to noise ratio. We developed a method analogous to that of (Zabeo et al., 2018), where the microtubule average volumes are aligned together in order to find their relative Φ rotation (Fig. S7 step 4). These Φ rotations were then applied to their respective particles and aligned to a common reference. SVA was subsequently performed with volumes binned 4, and 2 times. Only sporozoite data were aligned with unbinned volumes; the final step was to replace volumes reconstructed with the whole tilt-series by volumes reconstructed with ±24°. No rotation search was performed with the restricted tilt range. The resulting C1 EM map was anisotropic around the pseudosymmetry axis due to an uneven distribution of microtubule rotational orientations. To address this, particles were separated into classes by orientation and particles with the lowest cross correlation coefficients in the most abundant classes were removed. This reduced the number of particles from 24028 to 13263 in sporozoite and 8377 to 1851 in ookinete datasets.

#### Other microtubules

##### Gametocyte and merozoite singlet microtubules

Approximately 200 microtubules from 25 tomograms were processed individually, with random initial Φ rotations. Data were then manually classified by protofilament number, rotated to the same polarity and aligned with volumes binned 4 and 2 times. Helical parameters were measured directly using class averages with the exception of 13- and 15-protofilament classes where tubulin subunits were not resolved and published parameters were used. Helical symmetry was applied, followed by SVA alignment.

##### Gametocyte doublet microtubules

SPMT doublets were processed in a manner analogous to ookinete and sporozoite data, where average volumes were used to determine relative Φ rotations. SPMTs with different numbers of protofilaments in A and B tubules were combined together, resulting in an ensemble average volume.

### Half map generation

Particles were split into two halves (even and odd numbered particles). Random coordinate offsets of up to ±2 nm and random angular offsets of up to ±3° were added to each particle and the resulting parameters were used to re-generate a raw average for each half data set. Alignment of the two halves was then done independently with twice binned and then unbinned volumes, following the same procedure as the whole dataset. Fourier Shell Correlation was measured using Bsoft.

### Structure visualisation

EM maps and atomic models were visualised with UCSF (University of California San Francisco) Chimera (Pettersen et al., 2004) or UCSF ChimeraX (Pettersen et al., 2021). Computational sections were generated in IMOD.

### Multiple sequence alignment

ClustalOmega (Goujon et al., 2010; Sievers et al., 2011) was used to align TrxL1 protein sequences and JalView was used for visualisation (Waterhouse et al., 2009). Colours are based on ClustalX colouring.

### Tomogram segmentation for visualisation

Segmentation was performed manually in IMOD, using the drawing tools followed by linear interpolation. The resulting models were used to extract segmented volumes. SPMTs, ookinete conoid and sporozoite APR were “backplotted”: Average volumes were placed into 3D volumes using coordinates determined by SVA. SPMT and APR particles were spline fitted to smooth alignment errors for visualisation. Segmented and backplotted volumes were visualised using UCSF ChimeraX (Pettersen et al., 2021).

### Tomogram segmentation for IMC-MT distance and angular measurements

Tomograms were bandpass filtered using Bsoft. Segmentations of the filtered tomograms were guided by a tensor voting algorithm (Martinez-Sanchez et al., 2014). The parameters were optimised for each dataset, respectively. Clusters containing the inner membrane complex (IMC) segmentations were manually extracted by visual analysis. The clusters were then converted to a 3D point cloud and further processed using the Open3D library (Zhou et al., 2018). Statistical outlier analysis was used to remove excess noise from the segmentations. Subsequently, the DBSCAN algorithm was used to separate individual membrane sections and the outer side of the IMC was selected manually for subsequent distance measurements. The angle ΦIMC was measured between two vectors for each SPMT particle: A vector of the SPMT particle X axis and a vector from the particle centre to the nearest segmented IMC coordinate (roughly equivalent to IMC normal vector intersecting the SPMT particle). Particles outside segmented membrane patches were excluded. Measurements from individual microtubules were reduced to a median value for plotting and statistical analyses.

The requirement for a clear membrane density for automated segmentation substantially reduced the number of microtubules available for analysis. Thus, due to the rarity of SPMTs in merozoites, doublets in gametocytes and minus termini in ookinetes, a small number of tomograms were segmented manually in IMOD.

### Atomic model alignment and fitting

Initially, *T. gondii* SPMT model (PDB 7MIZ) was fitted to the *P. falciparum* sporozoite SPMT EM map as a rigid body in Chimera (Pettersen et al., 2004), the fit was then visualised in ChimeraX (Pettersen et al., 2021). The resulting fit was unambiguous with TgTrxL2 (which is not expressed in *Plasmodium* but is part of the *Toxoplasma* ILH) outside and the remaining protein chains inside the map. Due to the difference in ellipticity between the *P. falciparum* and *T. gondii* maps, protofilaments and TrxL1/SPM1 half arcs were fitted as separate units. Structural prediction of PfTrxL1 (https://alphafold.ebi.ac.uk/entry/Q8I2W0) (Jumper et al., 2021) was aligned with TgTrxL1 (PDB 7MIZ) using matchmaker (Meng et al., 2006) in ChimeraX.

### Protofilament angle measurements

Inter-protofilament angles were determined by measuring the angle between Cα atoms of the same pair of residues in neighbouring protofilaments. Median values of roughly 200 measurements were used.

| Parameter | Life cycle form |  |  |  |
| --- | --- | --- | --- | --- |
|  | Sporozoites | Ookinetes | Gametocytes | Merozoites |
| Number of tomograms used for SVA | 27 | 14 | 25 | 7 |
| Number of individual microtubules in final average | 407 | 159 | Total: 152<br>13: 10<br>14: 10<br>15: 14<br>16: 31<br>17: 62<br>18: 25 | Total: 12<br>Used in cfinal average: 8 |
| Total number of particles | 13263 | 1851 | 13: 4053<br>14: 11806<br>15: 10435<br>16: 21459<br>17: 42738<br>18: 15045 | 8532 |
| Pixel size (Å) | 3.335 | 6.67 (2x binning) | 6.67 (2x binning) | At bin2x: 9.05 |
| Symmetry<br>Helical:<br>Pitch (degrees),<br>helical rise (nm) | C1 | C1 | 13: 27.7, 0.94<br>14: 25.77, 0.87<br>15: 24, 1.08<br>16: 22.5, 0.99<br>17: 21.17, 0.93<br>18: 20, 0.90 | 27.7, 0.94 |
| EMDB accession number | EMDB Tbd | EMDB Tbd | EMDB Tbd | EMDB Tbd |

