## Supplemental Data for "Form follows function: Variable microtubule architecture in the malaria parasite"

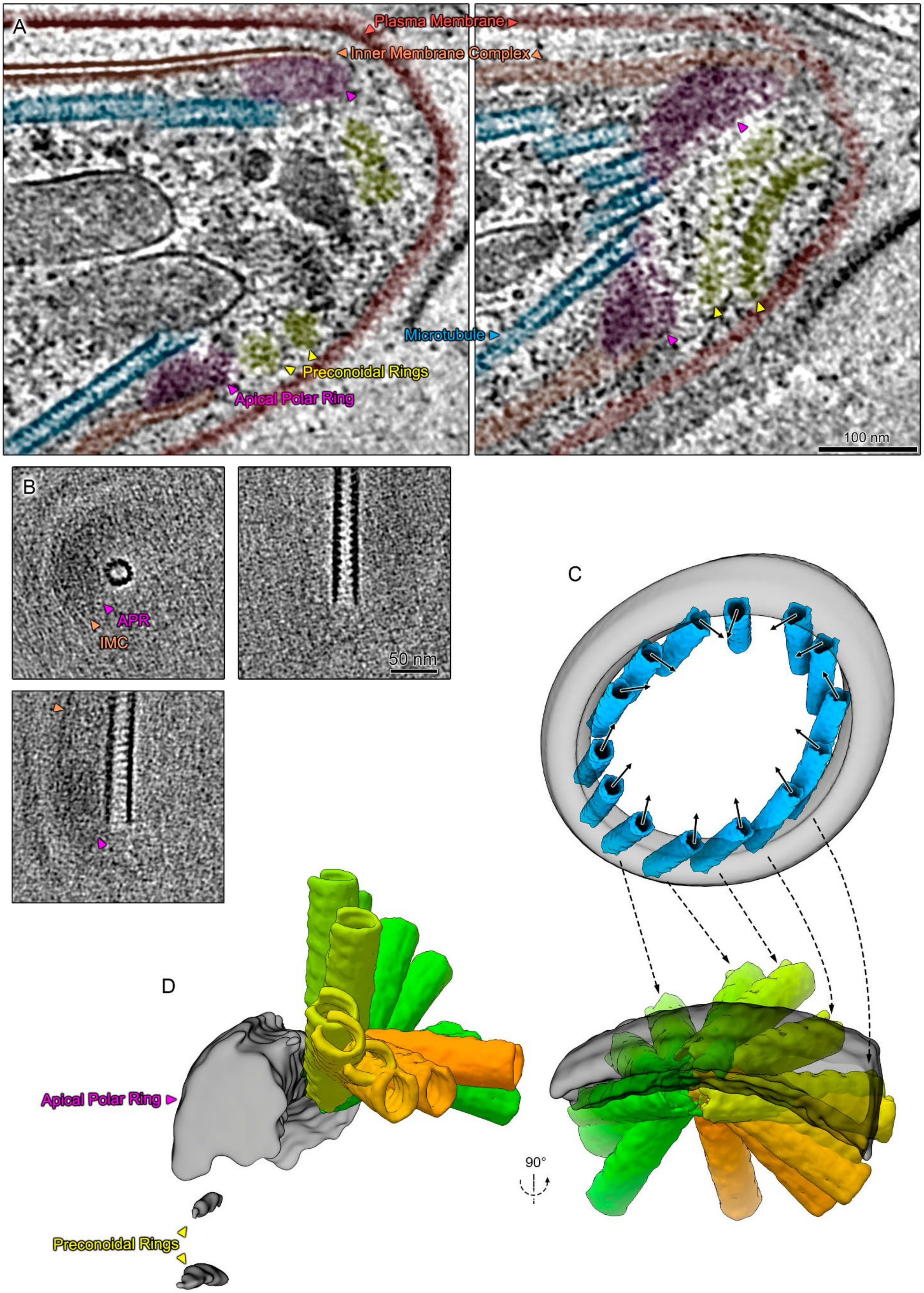
A
Plasma Membrane
Inner Membrane Complex
Microtubule
Preconoidal Rings
Apical Polar Ring
100 nm
B
APR
IMC
50 nm
C
D
Apical Polar Ring
Preconoidal Rings
90°

**Figure S1. Higher order organisation of subpellicular microtubules at the sporozoite apical pole. Related to Figure 2. A.** Two slices through the same tomogram of a sporozoite apex. Left: Slice through the centre of the APR and pre-conoidal rings. Right: slice through the edge of preconoidal rings and three SPMT minus ends. **B.** Orthogonal slices through an EM map of sporozoite SPMT minus (apical) ends. **C.** Computationally smoothed SVA model of an APR (grey) with copies of map shown in B (blue) placed at coordinates determined by SVA. Vectors representing the orientation of the seam relative to SPMT centre are represented with arrows. Dashed arrows indicate corresponding SPMTs, not all are indicated. **D.** Superposition of SPMT minus ends (orange to green) showing their orientation relative to the closest segment of APR (grey) and illustrating the large degree of freedom in binding. This was generated by shifting and rotating each SPMT volume to the coordinate system of the nearest APR coordinate determined by SVA. Weak densities for preconoidal rings can be seen in the APR EM map, showing they are very roughly aligned with the APR.

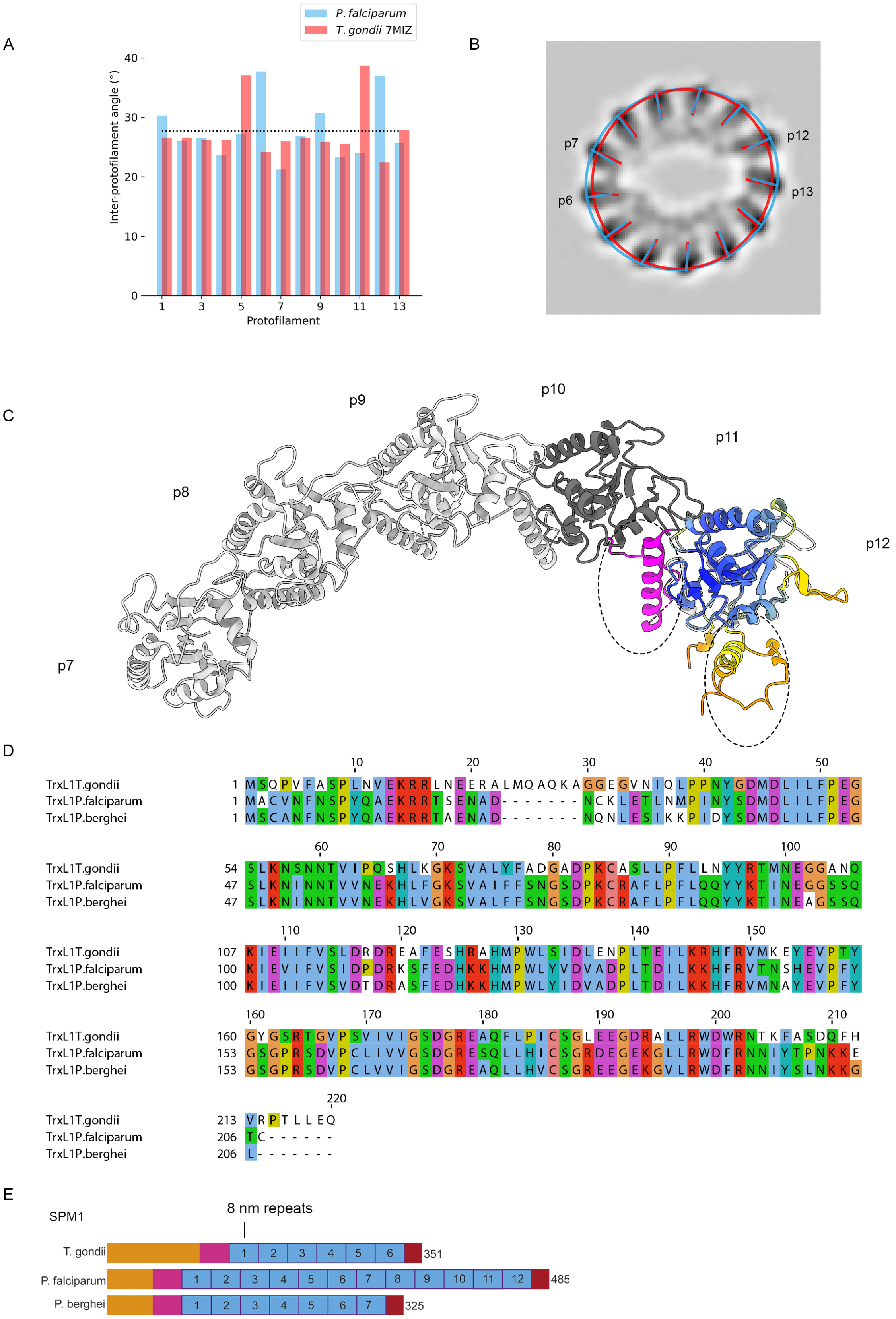
A
P. falciparum
T. gondii 7MIZ
Inter-protofilament angle (°)
0
10
20
30
40
1
3
5
7
9
11
13
Protofilament
B
p7
p6
p12
p13
C
p7
p8
p9
p10
p11
p12
D
10 20 30 40 50
TrxL1T.gondii 1 MSQPVFASPLNVEKRRLNEERALMQAQKAGGEGVNIQLPPNYGDMDLILFPEG
TrxL1P.falciparum 1 MACVNFNSPYQAEKRRTSENAD-------NCKLETLNMPINYSDMDLILFPEG
TrxL1P.berghei 1 MSCANFNSPYQAEKRRTAENAD-------NQNLESIKKPIDYSDMDLILFPEG
60 70 80 90 100
TrxL1T.gondii 54 SLKNSNNTVIPQSHLKGKSVALYFADGADPKCASLLPFLLNYYRTMNEGGANQ
TrxL1P.falciparum 47 SLKNINNTVVNEKHLFGKSVAIFFSNGSDPKCRAFLPFLQQYYKTINEGGSSQ
TrxL1P.berghei 47 SLKNINNTVVNEKHLVGKSVALFFSNGSDPKCRAFLPFLQQYYKTINEAGSSQ
110 120 130 140 150
TrxL1T.gondii 107 KIEIIFVSLDRDREAFESHRAHMPWLSIDLENPLTEILKRHFRVMKEYEVPTY
TrxL1P.falciparum 100 KIEVIFVSIDPDRKSFEDHKKHMPWLYVDVADPLTDILKKHFRVTNSHEVPFY
TrxL1P.berghei 100 KIEIIFVSVDTDRASFEDHKKHMPWLYIDVADPLTDILKKHFRVMNAYEVPFY
160 170 180 190 200 210
TrxL1T.gondii 160 GYGSRTGVPSVIVIGSDGREAQFLPICSGLEEGDRALLRWDWRNTKFASDQFH
TrxL1P.falciparum 153 GSGPRSDVPCLIVVGSDGRESQLLHICSGRDEGEKGLLRWDFRNNIYTPNKKE
TrxL1P.berghei 153 GSGPRSDVPCLVVIGSDGREAQLLHVCSGREEGEKGVLRWDFRNNIYSLNKKG
220
TrxL1T.gondii 213 VRPTLLEQ
TrxL1P.falciparum 206 TC------
TrxL1P.berghei 206 L-------
E
SPM1
8 nm repeats
T. gondii 1 2 3 4 5 6 351
P. falciparum 1 2 3 4 5 6 7 8 9 10 11 12 485
P. berghei 1 2 3 4 5 6 7 325

**Figure S2: The mosquito form microtubule is elliptical and has an interrupted luminal helix made up of the apicomplexan proteins TrxL1 and SPM1. Related to Figures 2 and 3. A.** Angle relative to protofilament n+1. Dotted line denotes the inter-protofilament angle of a theoretical 13 protofilament microtubule. Both the *Toxoplasma gondii* (red) structure and our mosquito form *Plasmodium* (red) structures have elliptical cross-sections. **B.** The average ellipticity of a *Plasmodium* (blue) and *T. gondii* (red) microtubule superimposed onto the *Plasmodium* EM map. The relative angle between protofilaments is depicted by an arbitrarily selected vector. **C.** Predicted structure of PfTrxL1 (coloured based on prediction confidence) aligned to one subunit of *T. gondii* TgTrxL1, showing a good agreement. One TgTrxL1 subunit is coloured in dark grey, with the first N-terminal helix in magenta. The largest difference between the predicted PfTrxL1 and experimentally determined TgTrxL1 is at this N-terminal helix, which is responsible for a large part of the subunit-subunit interface. This helix is not visible in the experimental map at the gaps in the ILH (next to protofilaments 6 and 12), likely due to flexibility. Interestingly, there is a 7 amino acid insertion in the first helix in *T. gondii* (L23-A29) relative to *P. falciparum*, but most of these residues are not involved in forming the interunit interface. **D.** Multiple sequence alignment of TrxL1 protein from *T. gondii* (TGGT1_115220) and its homologues in *P. falciparum* (PF3D7_0919300) and *P. berghei* (PBANKA_0820200). Note the 7 amino acid insertion in *T. gondii*. **E.** Domain architecture of the SPM1 protein from *T. gondii* (TGGT1_263520) and its homologues in *P. falciparum* (PF3D7_0909500) and *P. berghei* (PBANKA_0810700) . SPM1 proteins are formed of a series of 32 amino acid repeats (roughly 8 nm long) which have been duplicated in *Plasmodium* from 6 in *T. gondii* to 7 in *P. berghei* and 12 in *P. falciparum.* **E** is adapted from Tran et al., 2012.

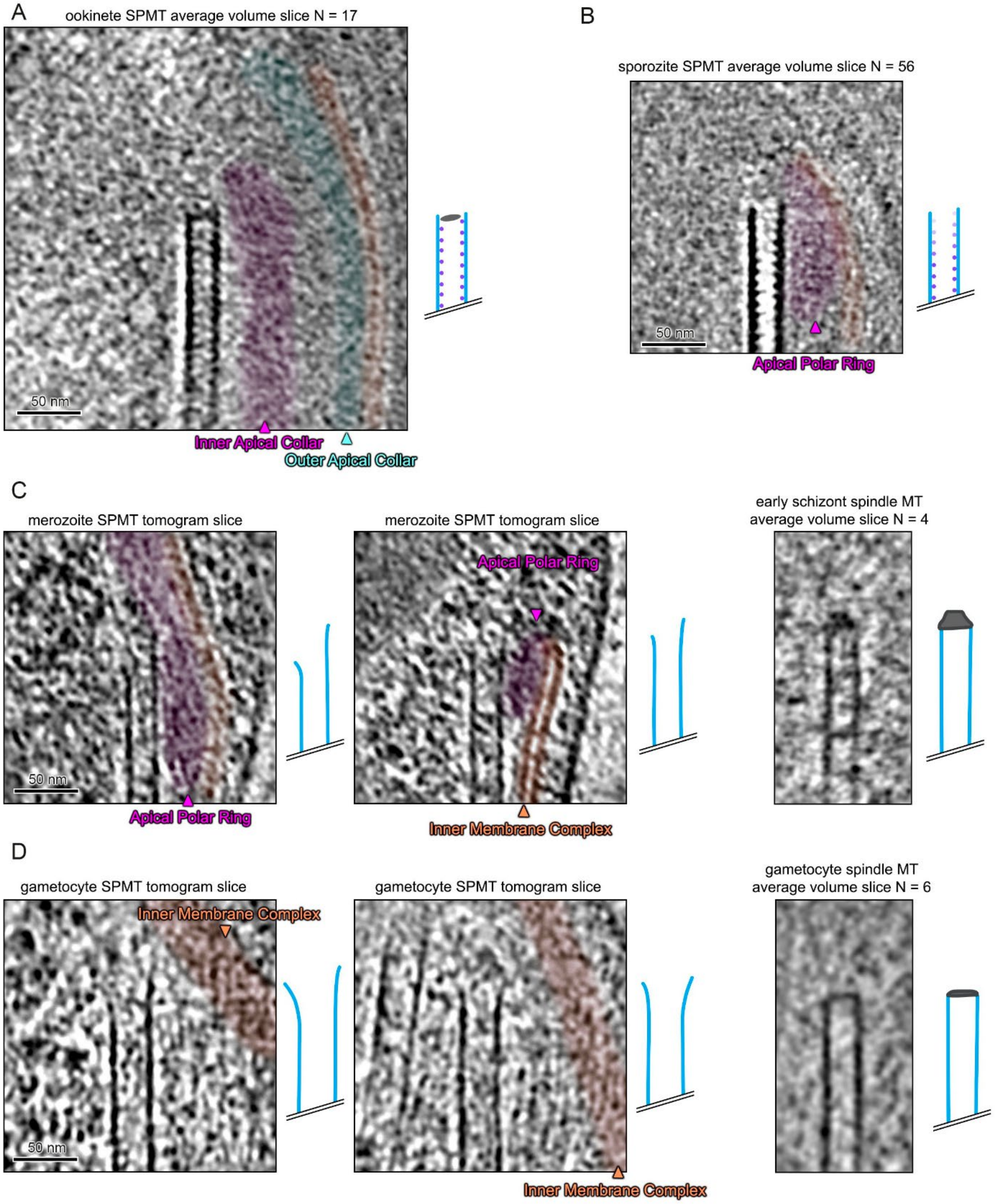


**Figure S3. Negative ends of SPMTs are uncapped, while nuclear spindle microtubules have a cap density consistent with γ-tubulin ring complex (γTuRC). Related to Figures 2,3,4,5.** The γ-tubulin ring complex (γTuRC) is a cone shaped complex consisting of 13 γ-tubulin subunits that presents binding sites to α- and β- tubulin. Cartoons to the right of greyscale images represent a model of the middle section through the respective microtubule minus ends. Number of particles included in SVA volumes are indicated above. IMC is highlighted in orange. **A.** Slice through an average volume of 17 negative ends of ookinete SPMTs. There was a hint of what could be a large protein complex in the terminus lumen, but it could also be due to a SVA alignment artefact combined with a low particle number. **B.** Slice through an average volume of 56 negative ends of microtubules from sporozoites. The tapering intensity of ILH towards the terminus could be due to progressively lower TrxL1 occupancy. **C**. Left: Slice through two example subvolumes of merozoite SPMT negative ends. Right: A slice through an average volume of 4 negative ends of nuclear spindle microtubules from schizonts. **D.** Slices through two examples of negative ends of gametocyte SPMTs and an average volume of 6 negative ends of nuclear spindle microtubules from gametocytes. This volume is an ensemble average containing primarily 15 protofilament microtubules.

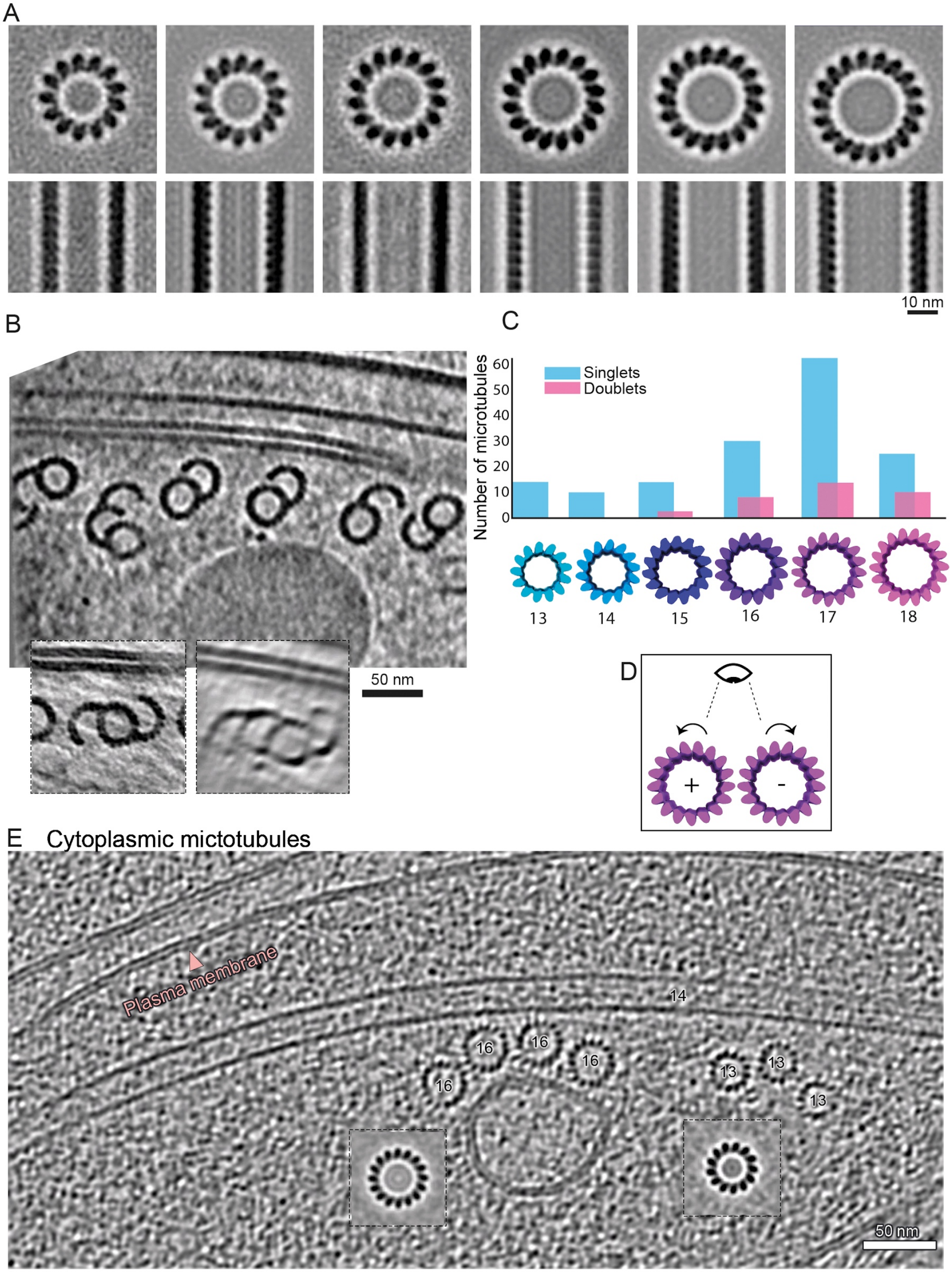

A
10 nm
B
50 nm
C
Number of microtubules
60
50
40
30
20
10
0
Singlets
Doublets
13
14
15
16
17
18
D
+
-
E Cytoplasmic mictotubules
Plasma membrane
14
16
16
16
16
13
13
13
50 nm

**Figure S4: Gametocyte SPMTs have random orientation and a range of protofilament numbers in singlets doublets, triplets and quadruplets**. **Related to figure 5. A.** Slices through two orthogonal axes of EM maps of 13 to 18 protofilament SPMTs. **B.** Example slice through a subvolume showing doublets and triplets with different geometries and orientations relative to the nearby IMC. Inset: Example slice through a subvolume showing a triplet and quadruplet SPMT (observed smearing is the result of missing tomographic information) **C.** Histogram of the protofilament number distribution in gametocyte singlets compared to doublet A tubules. **D.** Schematic showing how the polarity of a microtubule can be determined from the tilt direction of protofilaments. **E.** Slice through a subvolume of a stage III gametocyte cytoplasm showing cytoplasmic microtubules. In contrast to SPMTs, cytoplasmic microtubules appear clustered by protofilament number and polarity. Insets are examples of an average volumes of a single 13 protofilament microtubule and a single 16 protofilament microtubule from this tomogram.

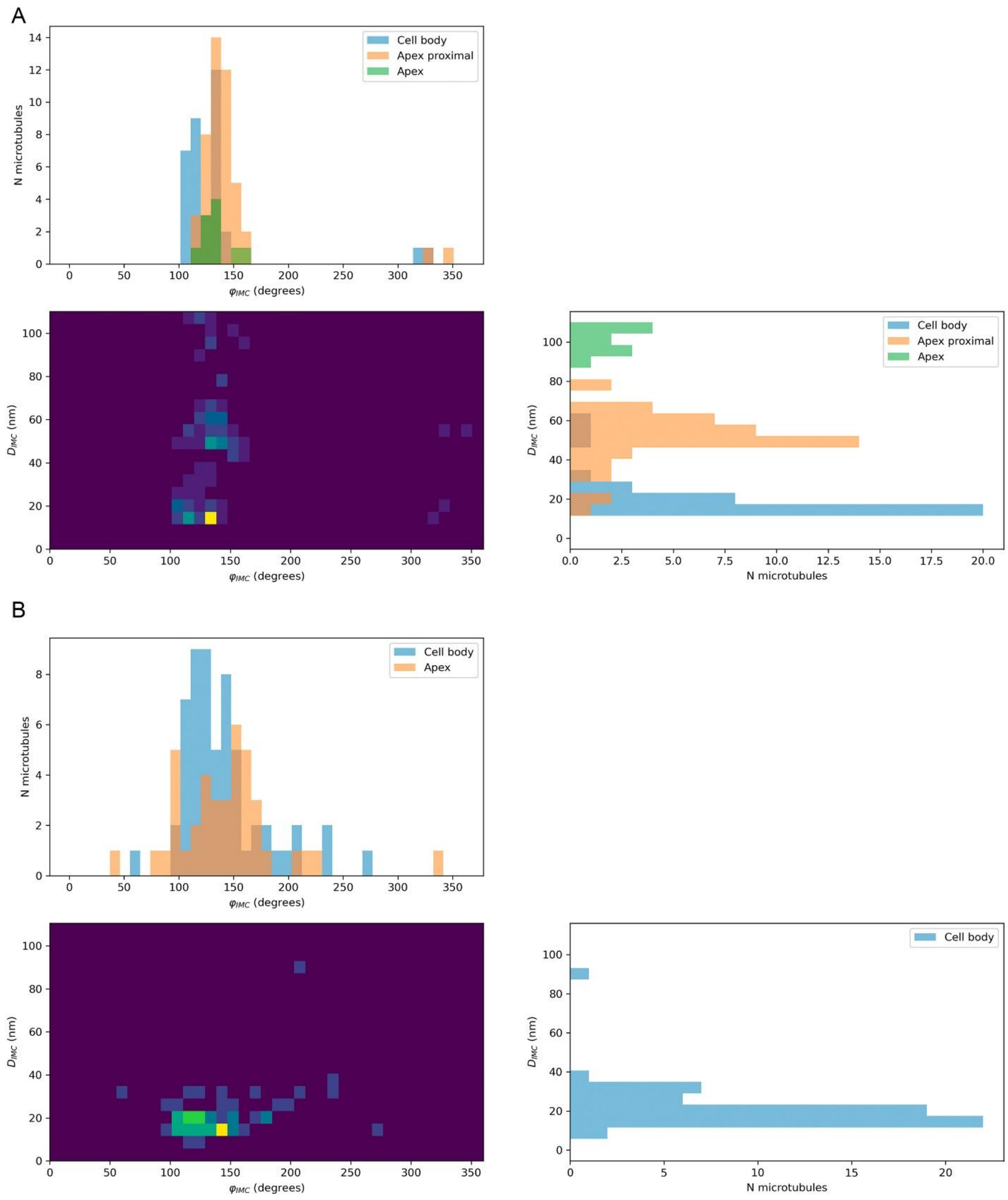


**Figure S5: Mosquito form microtubules have a conserved distance and angle to the IMC. Full dataset related to figure 6.** Linear 2D and 1D histogram representation of seam-IMC angle ($\varphi_{IMC}$) and SPMT-IMC distance ($d_{IMC}$) also shown in Fig 6. **A.** Ookinetes. **B.** Sporozoites.

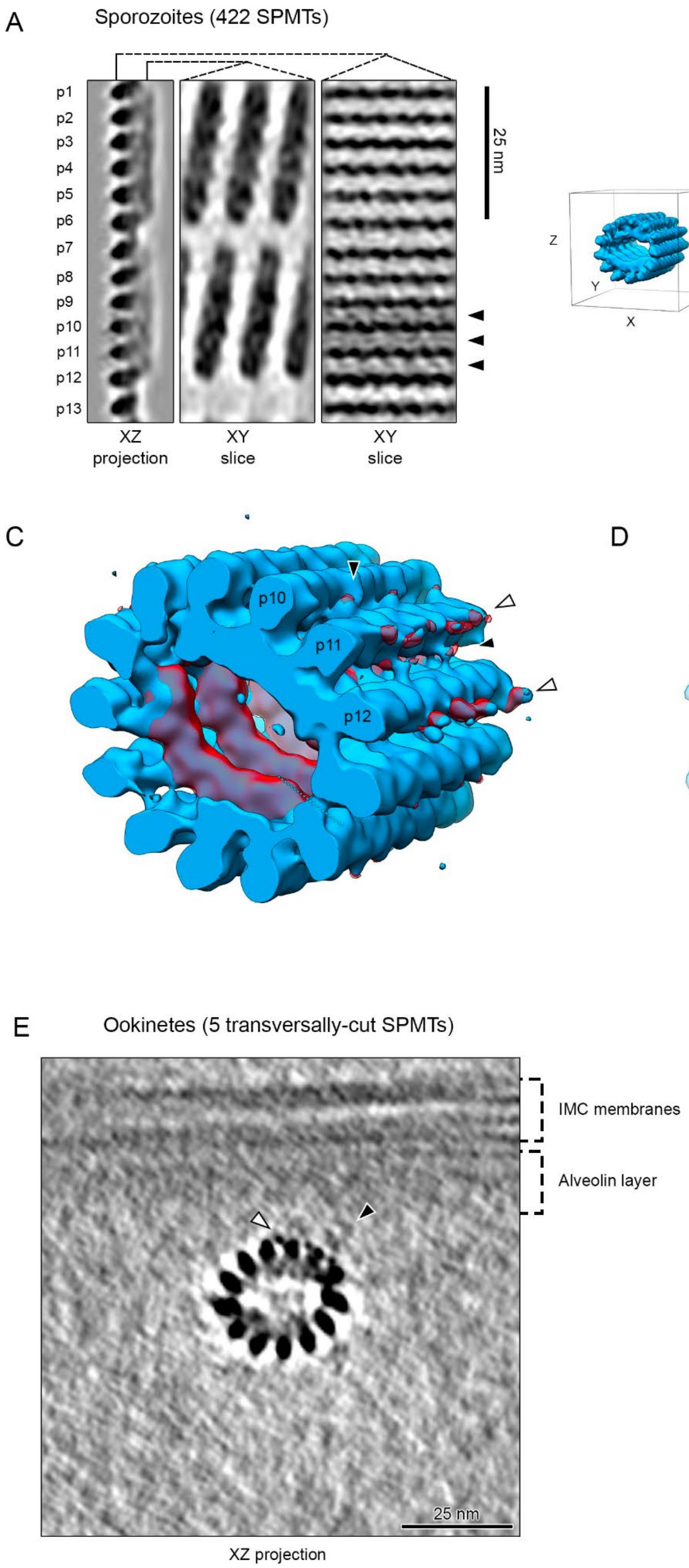


**Figure S6. Mosquito form microtubules have a radially-asymmetric decoration with an unknown protein between protofilaments 10, 11 and 12. Related to Figure 2 and 6. A.** The sporozoite SPMT EM map was “unwrapped” (radially projected) and shown sliced along different axes. Left: XZ projection equivalent to the original map projection in B, protofilaments numbers are indicated (equivalent to numbering in Fig. 3C). Middle: Section through the ILH layer, equivalent to a “panoramic” view from the inside of the SPMT. Right: Section glancing the outermost part of protofilaments. Black arrowheads indicate densities between protofilaments (also in B). **B**. Left: EM map isosurface with axes indicated to aid orientation. Right: Average projection through the EM map with contrast adjusted to highlight densities on protofilaments 10 and 11 (white arrowheads, also in C). **C**. Isosurface of SPMT EM map (blue) overlapped with a difference map (red) resulting from subtracting a simulated tubulin density map. **D.** Isosurface of three adjacent protofilaments with fitted tubulin model (7MIZ). Black dashed circles highlight extra densities between protofilaments. Note that these densities form a bridge between two α-tubulin subunits (between protofilaments 10 and 11) and two β-tubulin subunits (between protofilaments 11 and 12). **E**. Average projection through EM map generated by averaging 5 ookinete SPMTs that were cut at an almost perfect transversal orientation (i.e. their long axis was aligned with the electron beam at 0° stage tilt). The asymmetric densities seen in the sporozoite SPMT EM map (B) are clear, as well as the membrane and alveolin layers of the IMC. This suggests that the densities emanating from protofilaments 11 and 12 are in fact parts of the SPMT-IMC link. However, most SPMTs are aligned with protofilaments 8 being closest to the IMC (i.e. the average orientation of the IMC is tilted ~ 45° to the right), indicating that there might be additional links to protofilaments 6 to 10 stabilising this orientation.

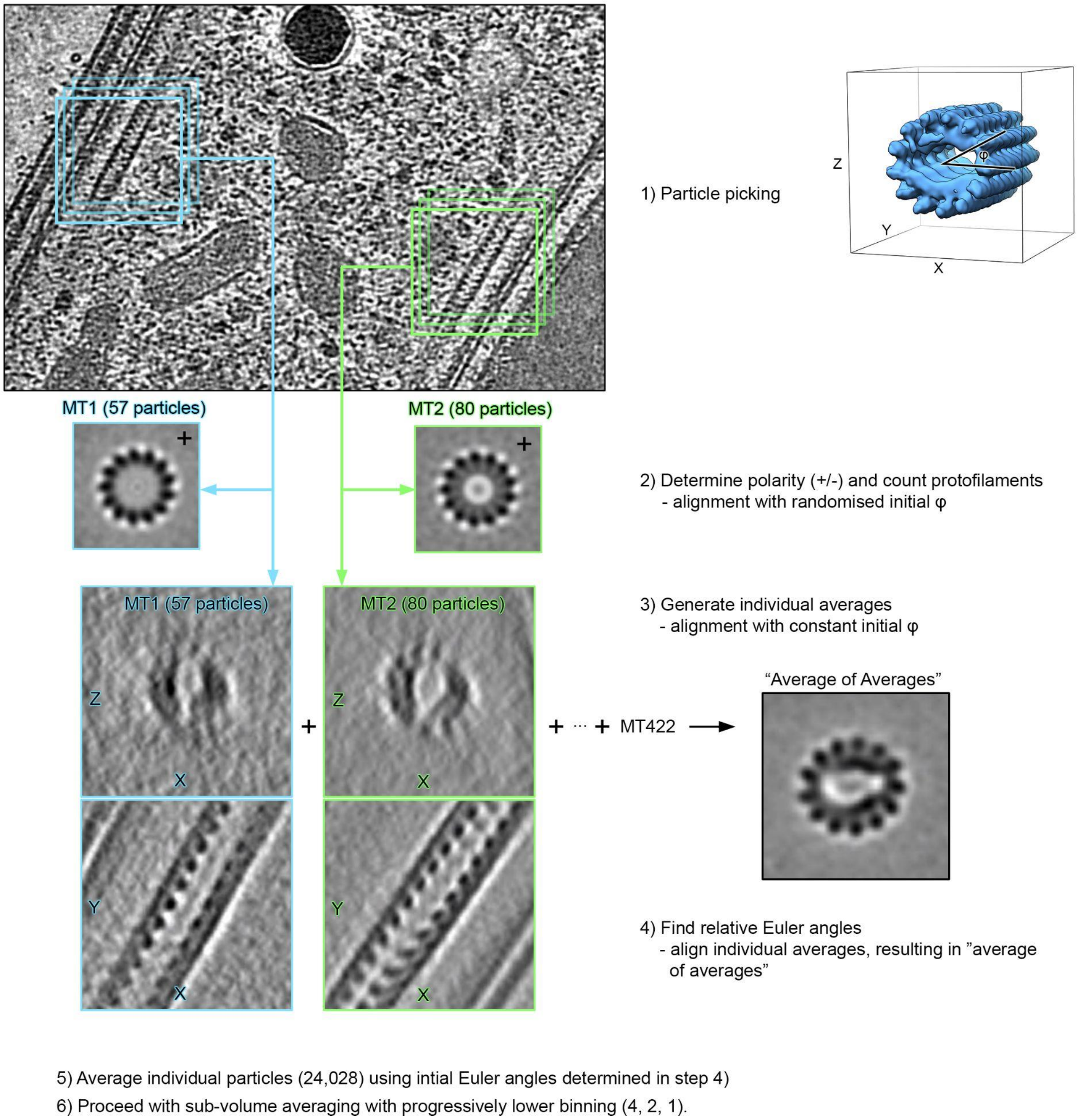


**Figure S7: Schematic workflow for subvolume averaging of mosquito form SPMTs using sporozoite data as an example. Related to Methods.** 1) Microtubules were manually picked by tracing their centres, then interpolated to generate regularly spaced particles. 2) Each microtubule was processed individually with randomised initial φ angles (angle around pseudosymmetry axis) and these average volumes were used to determine the polarity of each microtubule (e.g. Fig. S5 D). 3) Each microtubule was averaged individually with constant initial φ angles, resulting in C1 average volumes. The missing tomographic information ("missing wedge") is apparent in these. 4) Individual average volumes from step 3) were now aligned together to make an "average of averages", thereby determining their relative φ rotation. 5) The newly determined φ angles were applied to individual particles. 6) SVA was performed at progressively lower binning until a final unbinned average was produced.

| Tomo# | 13pf - | 13pf + | 14pf - | 14pf + | 15pf - | 15pf + | 16pf - | 16pf + | 17pf - | 17pf + | 18pf - | 18pf + | Total - | Total + |
|---|---|---|---|---|---|---|---|---|---|---|---|---|---|---|
| 1 | 0 | 0 | 0 | 1 | 0 | 0 | 1 | 1 | 1 | 1 | 0 | 0 | **2** | **3** |
| 2 | 0 | 0 | 0 | 0 | 3 | 0 | 2 | 3 | 0 | 1 | 0 | 0 | **5** | **4** |
| 3 | 0 | 0 | 1 | 0 | 0 | 3 | 0 | 0 | 2 | 0 | 0 | 2 | **3** | **5** |
| 4 | 0 | 1 | 1 | 1 | 0 | 0 | 3 | 0 | 3 | 5 | 1 | 3 | **8** | **10** |
| 5 | 0 | 0 | 1 | 0 | 0 | 0 | 2 | 3 | 0 | 6 | 1 | 3 | **4** | **12** |
| 6 | 0 | 1 | 0 | 0 | 1 | 3 | 1 | 1 | 0 | 1 | 0 | 0 | **2** | **6** |
| 7 | 0 | 0 | 0 | 0 | 0 | 0 | 0 | 0 | 0 | 1 | 1 | 3 | **1** | **4** |
| 8 | 0 | 0 | 0 | 0 | 1 | 0 | 0 | 0 | 3 | 1 | 1 | 0 | **5** | **1** |
| 9 | 0 | 0 | 0 | 0 | 0 | 0 | 0 | 0 | 4 | 2 | 1 | 1 | **5** | **3** |
| 10 | 0 | 1 | 0 | 0 | 0 | 0 | 1 | 0 | 2 | 2 | 0 | 0 | **3** | **3** |
| 11 | 0 | 0 | 0 | 0 | 0 | 0 | 0 | 1 | 1 | 2 | 0 | 0 | **1** | **3** |
| 12 | 0 | 0 | 0 | 0 | 0 | 0 | 0 | 0 | 2 | 1 | 0 | 0 | **2** | **1** |
| 13 | 0 | 4 | 0 | 0 | 0 | 0 | 0 | 1 | 2 | 2 | 0 | 0 | **2** | **7** |
| 14 | 1 | 0 | 1 | 0 | 1 | 0 | 0 | 0 | 1 | 1 | 0 | 0 | **4** | **1** |
| 15 | 1 | 1 | 0 | 0 | 1 | 0 | 0 | 0 | 0 | 2 | 0 | 0 | **2** | **3** |
| 16 | 0 | 0 | 0 | 0 | 0 | 1 | 0 | 2 | 0 | 1 | 0 | 1 | **0** | **5** |
| 17 | 0 | 0 | 1 | 0 | 0 | 0 | 2 | 1 | 3 | 4 | 2 | 2 | **8** | **7** |
| 18 | 0 | 0 | 1 | 1 | 0 | 0 | 1 | 0 | 4 | 1 | 1 | 0 | **7** | **2** |
| 19 | 0 | 0 | 0 | 0 | 0 | 0 | 0 | 0 | 0 | 0 | 1 | 0 | **1** | **0** |
| 20 | 0 | 0 | 0 | 0 | 0 | 0 | 0 | 0 | 0 | 0 | 1 | 0 | **1** | **0** |
| 21 | 4 | 0 | 0 | 1 | 0 | 0 | 4 | 0 | 0 | 0 | 0 | 0 | **8** | **1** |
| **Total** | **6** | **8** | **6** | **4** | **7** | **7** | **17** | **13** | **28** | **34** | **10** | **15** | 74 | 81 |

**Table S1: Polarities of gametocyte SPMTs in individual tomograms separated by protofilament number.**
